# A bare host cell membrane with minimal glycocalyx is an optimal surface for targeting by virulence-primed *Salmonella* Typhimurium

**DOI:** 10.64898/2026.05.11.724231

**Authors:** Petra Geiser, Johanna Westman, Merve Ceylan, Willem Bosman, Christopher von Beek, Per Artursson, Lena Kjellén, Thaher Pelaseyed, Mikael E. Sellin

## Abstract

Gut pathogens such as *Salmonella enterica* serovar Typhimurium target intestinal epithelial cells for adhesion and type-3-secretion-system-dependent invasion, while also invading multiple other cell types as the infection progresses. Mechanistic studies have uncovered virulence factors involved in this process, but host cell determinants affecting *Salmonella* cell surface targeting remain less deeply explored. Furthermore, cell surface protein expression and glycosylation patterns differ dramatically between epithelial and blood-derived cell types, and even across maturation states of the same cell type. Here, we explored bottom-up reconstruction of the host cell surface, using simplistic suspension-growing K562 cells, to determine the contribution of individual cell surface constituents during *Salmonella* targeting. Combined with flow cytometry and a stringently tunable gene expression system, this model enabled high-throughput analysis and combined genetic manipulations in both pathogen and host cells. Transcriptomic and proteomic data along with lectin characterization revealed minimal K562 surface glycosylation at baseline. Chemical manipulations substantiated the role of cell membrane cholesterol in promoting *Salmonella* targeting, whereas ectopic expression of glycoproteins such as transmembrane mucins introduced a size-dependent steric barrier towards invading bacteria. Strikingly, even established glycoprotein receptors for *Salmonella* adhesins hampered rather than promoted invasion, suggesting that adhesins are required to overcome cellular glycocalyces *in vivo*, while a bare host cell membrane would be the pathogen’s preferred interaction surface.

## INTRODUCTION

Gut bacterial pathogens including *Salmonella enterica* serovar Typhimurium (*Salmonella*) and close relatives are ingested along with contaminated food or water, colonize the intestinal lumen and target the epithelial surface for invasion. *Salmonella* targeting of intestinal epithelial cells (IECs) requires flagellar motility for penetration of the MUC2-based mucus layer secreted by intestinal goblet cells (1). Upon reaching the epithelial surface, *Salmonella* engages in flagellar-mediated near surface swimming, which allows the pathogen to scan the surface for suitable binding and invasion sites (2). Epithelial topology and *Salmonella* adhesin-mediated transient interactions with cell surface structures can mediate stopping and reversible adhesion (2–4). Such events allow for insertion of the syringe-like type-3-secretion-system-1 (T3SS-1) into the host cell membrane, resulting in irreversible docking (3) and translocation of effector proteins to trigger uptake (5, 6). Effector-induced remodeling of the host cell actin cytoskeleton entraps *Salmonella* in an endocytic vacuole, which the bacterium transforms into a specialized intracellular niche through a second T3SS encoded on *Salmonella* pathogenicity island 2 (SPI2) (7). A subpopulation of intracellular *Salmonella* also escapes the vacuole to hyper-replicate in the cytosol (8). In addition to absorptive IECs, *Salmonella* has been shown capable of entering a wide variety of other cell types *in vivo* and/or in culture, such as M cells, goblet cells, monocytic and lymphocytic cells, fibroblasts and mast cells (9–13).

*Salmonella* expresses multiple fimbrial and non-fimbrial adhesins that interact with a plethora of host cell surface structures to establish adhesion and invasion in different contexts (10, 14–20). Many adhesins have been shown to target glycan-based structures such as heavily glycosylated transmembrane mucins or proteoglycans (10, 14–17, 20), or in some cases dedicated surface receptors (18). In addition, bacterial lipopolysaccharide has been described to bind host glycans (21, 22), flagella may interact with the host cell membrane to promote adhesion (23, 24), and insertion of the T3SS-1 tip proteins SipB and SipC into the membrane mediates intimate host cell attachment (25, 26). Despite the pathogen’s large repertoire of adhesins and other surface structures, and the fact that cooperative action among some of them has been described (3, 27), adhesin requirements for targeting of different cellular surfaces vary. For instance, invasion of polarized epithelial cells is impaired in the absence of the non-fimbrial giant adhesin SiiE encoded on SPI4, while targeting of non-polarized cells does not require SiiE (28, 29). Furthermore, the different stages of host cell targeting are determined by various, and oftentimes correlated, host cell-intrinsic factors such as cell morphology, membrane composition, cell cycle state and expression of non-glycosylated and glycosylated cell surface structures. There have been efforts to systematically assess these cell-intrinsic vulnerabilities (30), but the impact of specific host cell features remains partially elusive. When it comes to cell surface glycostructures such as proteoglycans and transmembrane mucins abundantly expressed in the IEC apical brush border *in vivo*, these structures have been implied as both a steric barrier towards commensal and pathogenic bacteria (31–41) and as receptors for bacterial adhesins, which would promote host cell targeting (10, 14–17, 20–22, 40, 42, 43). Which of these aspects is decisive during host cell colonization remains to be determined.

Here, we developed a simplistic cell line-based high-throughput infection model suitable to answer these questions and systematically assess how surface characteristics affect targeting by *Salmonella*. Due to the short doubling time of ∼18-24 h, available genetic tools, and their suspension-growing properties, K562 cells represent a scalable model system compatible with facile high-throughput analyses. The main cell type targeted by *Salmonella* invasion *in vivo*, IECs, are covered in a dense and complex glycocalyx where it is difficult to pinpoint the contribution of individual protein and glycan structures in either promoting or blocking bacterial targeting. As the majority of these structures are absent in non-epithelial K562 cells, this model is ideally suited for bottom-up reconstruction of host cell surface constituents. The availability of a tightly controllable expression system in this cell line (44–46) allowed us to study the role of individual, ectopically expressed surface glycoproteins during host cell targeting. Collectively, our results indicate that T3SS-1-bearing *Salmonella* preferentially targets a bare, cholesterol-rich membrane, while cell surface glycostructures, as well as bacterial appendages, generally form a size-dependent barrier to invasion. Hence, we propose that the large repertoire of adhesins is the result of *Salmonella* being forced to overcome complex host cell surface glycoprotein barriers, whereas a bare plasma membrane would constitute the optimal targeting surface from the pathogen’s point of view.

## RESULTS

### A versatile high-throughput *Salmonella* infection model using suspension-grown K562 cells

K562 cells offer an attractive opportunity as a simple, scalable and high-throughput infection model compatible with flow cytometry analysis. To that end, we first verified if K562 cells were permissive to *Salmonella* infection, by adapting the Gentamicin protection assay. Intracellular *Salmonella* were detected by fluorescence microscopy via an intracellular vacuolar, SPI2-driven reporter (p*ssaG*-GFP; (47, 48). GFP-positive bacteria localized to the perinuclear area, where the SCV is expected to be found (Fig 1A; (7)). Next, we assessed whether *Salmonella* invasion and establishment of a vacuolar niche could be analyzed by flow cytometry. Parallel invasion assays in K562 and a well-established infection model, adherent HeLa cells (7), were performed and the percentage of infected, *ssaG*-GFP-positive cells quantified (Fig 1B). The results indicated that *Salmonella* invasion of K562 cells increased with multiplicity of infection (MOI) broadly similar to invasion in HeLa cells, albeit with modestly lower efficiency (Fig 1C). Furthermore, infections with a panel of T3SS-1 effector mutants (Fig 1D) revealed that invasion in K562 cells required the same set of effectors, the ruffle-inducers SopB, SopE and SopE2, as previously established for HeLa cells, but with an additional contribution of SipA, an actin filament stabilizer previously shown to be critical for invasion of non-transformed IECs (29, 33). To assess intracellular replication, K562 and HeLa cells were infected with reporter-less *Salmonella* and intracellular bacteria extracted and plated for colony-forming units (CFU) at 2 h and 18 h post infection (p.i.). This demonstrated *Salmonella* replication within both K562 and HeLa (Fig 1E).

**Figure 1.**
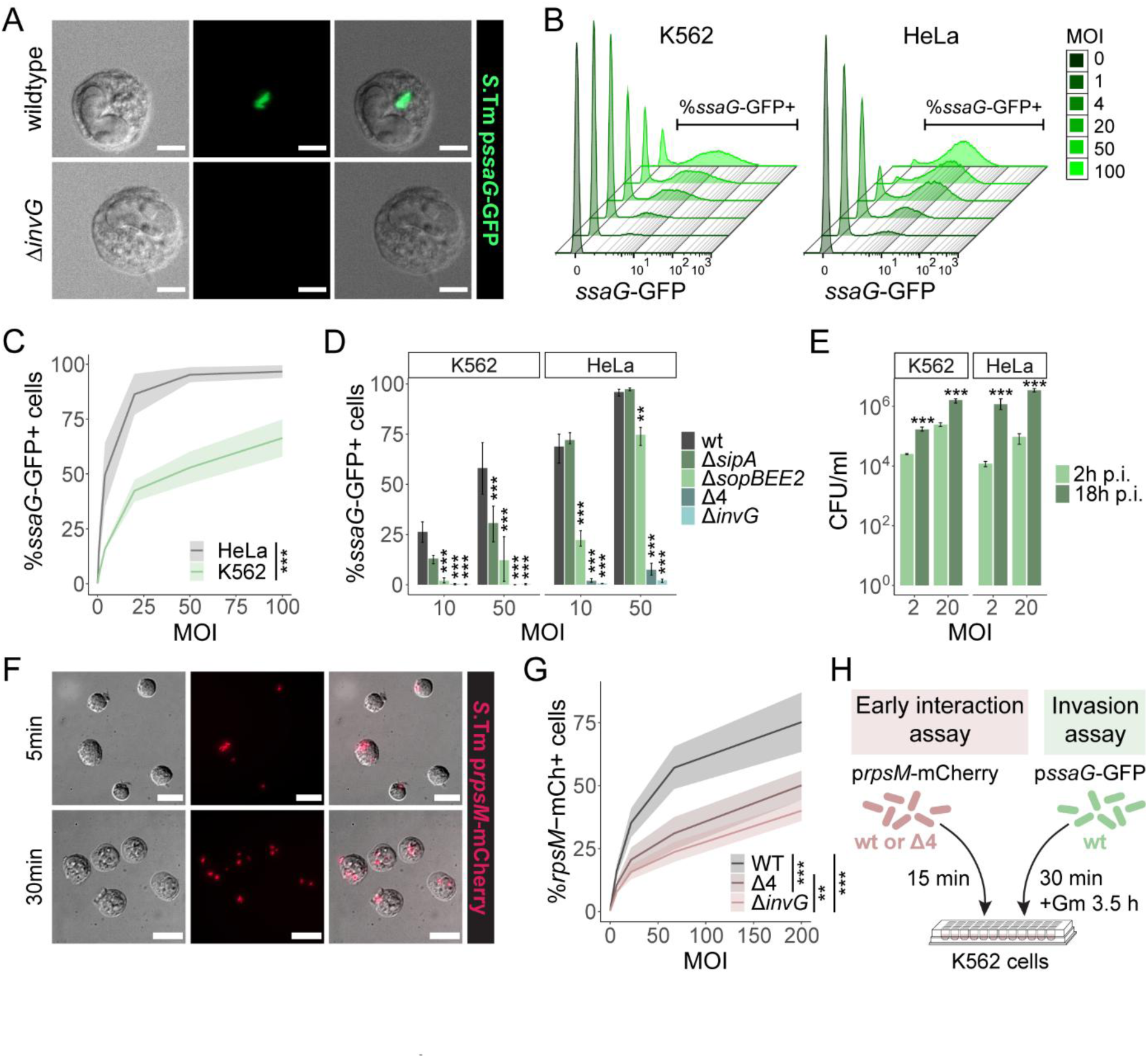
Establishment of a high-throughput *Salmonella* infection model. (A) K562 cells were infected with the indicated *Salmonella* (*S*.Tm) strain harboring the intracellular vacuolar p*ssaG*-GFP reporter for 30 min, followed by Gentamicin treatment and reporter maturation for an additional 3.5 h. Overlays of a single-plane image for the differential interference contrast (DIC) channel and maximum intensity projections for the GFP channel are shown. Scale bars: 10 µm. (B-D) K562 and HeLa cells were infected in parallel as in A and (B) analyzed by flow cytometry to determine (C) the percentage of infected (*ssaG*-GFP-positive) cells. (D) Infection with a panel of *Salmonella* mutants reveals the T3SS-1 effector requirements for invasion of K562 and HeLa cells. (E) Intracellular *Salmonella* replication in K562 and HeLa cells was assessed by CFU (colony forming units) plating at 2 h and 18 h p.i. in a Gentamicin protection assay. Data is plotted as CFU/well, with a well harboring ∼750’000 cells. (F-G) Establishment of an early interaction assay with constitutively fluorescent *Salmonella* p*rpsM*-mCherry strains inoculated with K562 cells. (F) Fluorescence microscopy confirms specific detection of cell-associated bacteria without contamination by unbound bacteria. Scale bars: 10 µm. (G) Parallel infection of K562 cells with *Salmonella* wt, Δ4 and Δ*invG* indicates that both adhesion and early invasion are quantified, and reveals a role for T3SS-1 during stable binding. (H) Schematic of the experimental setup for early interaction and invasion assays in round-bottom 96-well plates compatible with flow cytometry. Data is plotted as mean + range of 3 replicates (D, E); or as mean + standard deviation (sd) of ≥ 4 replicates (C, G). Statistical analysis was performed by 2-way ANOVA (C-E, G) with Tukey’s honestly significant difference (TukeyHSD) post-hoc test (D, E, G). **, p < 0.01; ***, p < 0.001, MFI; median fluorescence intensity.

Scaling up infections to a 96-well plate format yielded comparable *Salmonella* invasion efficiencies (Fig S1A) and allowed for parallel infections with multiple T3SS-1 effector mutants over multiple MOIs in a single experiment (Fig S1B; compare to Fig 1D). Since a subpopulation of *Salmonella* is known to inhabit a cytosolic niche in epithelial cells (8), we quantified vacuolar and cytosolic *Salmonella* frequencies, relying on parallel infections with *Salmonella* harboring either p*ssaG*-GFP or a cytosol-specific p*uhpT*-GFP reporter (4, 49). No more than 7% of infected K562 cells harbored cytosolic bacteria, whereas the population containing vacuolar *Salmonella* reached ∼50% at the highest MOIs (Fig S1C). This corresponds to the ratio of cytosolic to vacuolar *Salmonella* documented in other cell lines (8, 50, 51) and means that the p*ssaG*-GFP reporter strain can be confidently used to quantify a major fraction of the colonization events in K562.

To study initial *Salmonella* targeting of the host cell surface, we established an additional assay in K562 cells based on a constitutive bacterial reporter (p*rpsM*-mCherry; (52)). The use of Cytochalasin-D to block actin-mediated bacterial uptake during *Salmonella* wt infection resulted in low numbers of cell-associated bacteria after washing (Fig S1D). This suggests that adhering *Salmonella* can detach during centrifugation and washing, and challenges the concept of T3SS-1-mediated irreversible docking (3). Instead, a reduction in infection time provided an ‘early interaction assay’ (Fig S1E) and fluorescence microscopy validated specific quantification of cell-associated bacteria (Fig 1F). *Salmonella* adhered to K562 cells already by 5 min, and following 30 min interaction, both surface-adherent and clearly intracellular bacteria could be observed (Fig 1F). 15 min was chosen as an optimized infection time for the early interaction assay. Parallel infections with *Salmonella* wt and two mutants lacking either the entire T3SS-1 (Δ*invG*, (53)), or its four main effectors, SipA, SopB, SopE and SopE2 (Δ4, (54)), indicated that this assay could differentiate between exclusively adhesion versus adhesion and early invasion, as host cell association for the non-invasive Δ4 strain was reduced by ∼30-40% relative to the wt (Fig 1G). A further reduction for the Δ*invG* mutant confirmed a role for the T3SS-1 apparatus itself in host cell surface binding (Fig 1G), as previously noted in adherent cell lines (3, 26). In conclusion, we established a scalable, high-throughput, flow cytometry-based *Salmonella* infection model in K562 cells, which is well-suited to assess (i) early interaction with the host cell surface, as well as (ii) invasion and vacuolar colonization, across a range of conditions (Fig 1H).

### Membrane cholesterol levels positively affect *Salmonella* targeting of actively growing as well as mitotically blocked K562 cells

Previous studies have consistently shown that *Salmonella* preferentially targets mitotic cells in culture (2, 55, 56). However, alternative mechanisms have been proposed. On one hand, increased cholesterol during mitosis is thought to increase docking via the T3SS-1 translocon protein SipB (25, 26, 56). Alternatively, the basis for preferential targeting might be purely physical, as adherent cells round up during mitosis, therefore representing topological obstacles that can increase the frequency of *Salmonella* stopping and docking during near-surface swimming (2). Due to their suspension-growing properties, the morphology of K562 is constant throughout the cell cycle, making this model ideally suited to disentangle this question. To that end, K562 cells were treated with Taxol for 10 min as a control, or overnight (o/n, ∼18 h) to block formation of the mitotic spindle. This resulted in cell cycle arrest and an enrichment of the mitotic population without affecting cell viability (Fig 2A-B, S2A-B). Filipin staining indicated increased cholesterol levels in mitosis-arrested cultures (Fig 2C). Still, an early interaction assay refuted preferential targeting of mitotic cells by *Salmonella* in this experimental setup (Fig 2D). Interestingly, assessment of cholesterol levels in conjunction with the early interaction assay consistently showed increased Filipin intensities for the *Salmonella*-positive cell population relative to the negative subpopulation (Fig 2E). This suggests that high cholesterol content is a predictor for *Salmonella* targeting of the host cell surface regardless of cell cycle phase.

**Figure 2.**
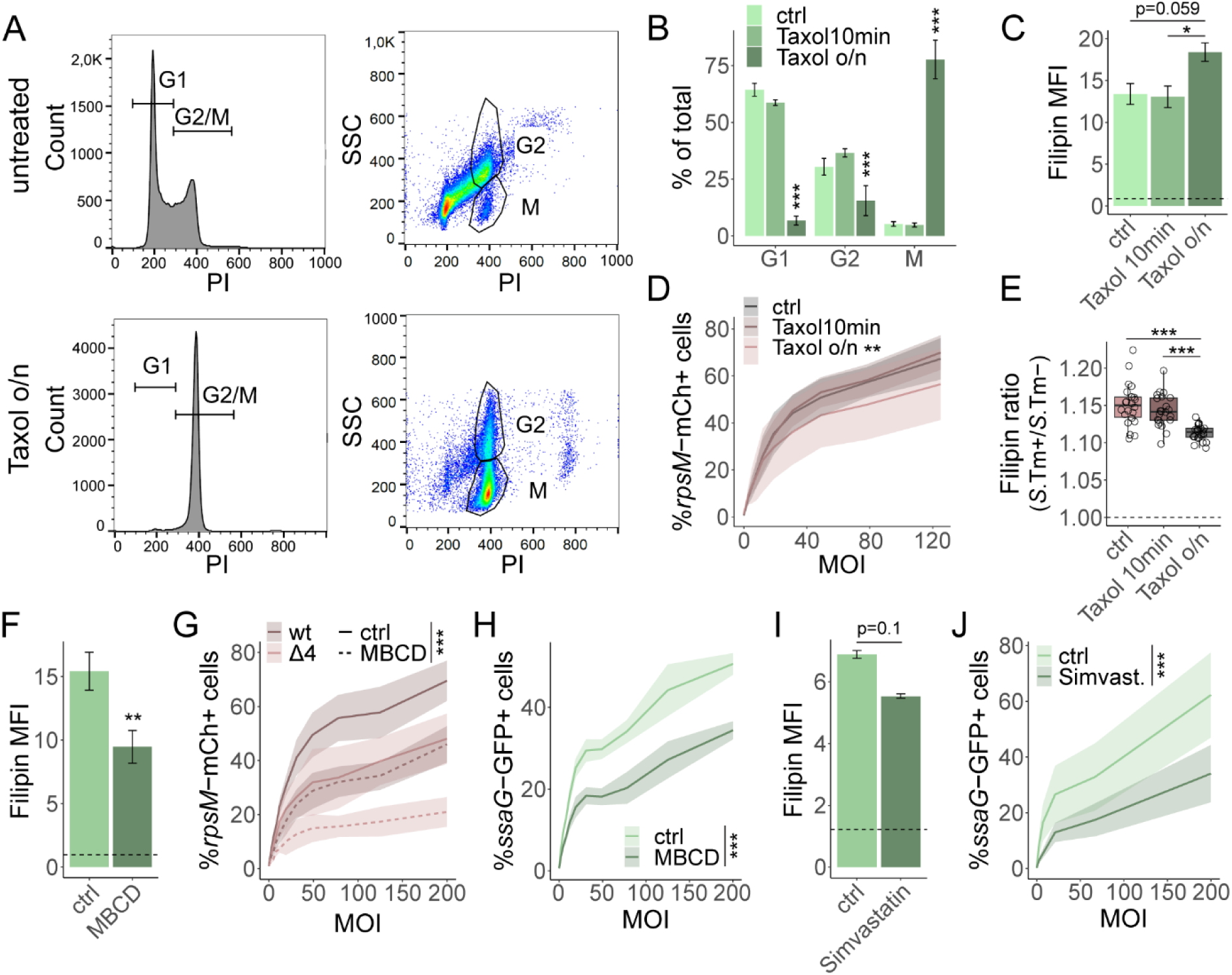
Cholesterol levels impact *Salmonella* host cell targeting. (A-B) Overnight (o/n) Taxol treatment arrests K562 cells in mitosis. (A) Untreated (top) or o/n Taxol-treated (bottom) K562 cells were permeabilized with 0.1% Triton-X-100 and stained with propidium iodide (PI) to distinguish between G1 and G2/M phase cells based on the amount of DNA per cell (left panels), with G2/M cells displaying twice the intensity of G1 cells. G2 and M cells can be distinguished based on their side scatter (SSC), as granularity decreases upon nuclear membrane degradation during mitosis. (B) The percentages of cells assigned to each cycle phase show an increase in the mitotic population for o/n Taxol-treated cells. Data is plotted as mean + sd of 4 replicates and statistical analysis was performed by 2-way ANOVA and TukeyHSD. (C) Filipin staining indicates an increase in cell surface cholesterol levels in mitotic-arrested cells. Data is plotted as mean + sd of 4 replicates and statistical analysis was performed by Kruskal-Wallis test with Dunn’s post-hoc test and Benjamini-Hochberg adjustment for multiple comparisons. (D) No preferential *Salmonella* targeting of mitotic-arrested cells was observed in an early interaction assay. Data is plotted as mean + sd of ≥ 11 replicates and statistical analysis was performed by 2-way ANOVA and TukeyHSD. (E) Subsequent Filipin staining of infected cells and calculation of the Filipin MFI ratio in *Salmonella* p*rpsM*-mCherry-positive over -negative cells reveals that cholesterol-enriched cells are preferentially targeted across all conditions. Pooled data for all MOIs >0 from 2 replicates is plotted. In boxplots, the height of the boxes represents the interquartile range (IQR), whereas the horizontal line depicts the median. Whiskers extend to the most extreme data point within 1.5x IQR. All data points are indicated as circles. (F-H) Methyl-β-cyclodextrin (MBCD) treatment for 30 min reduces (F) cell surface cholesterol levels as determined by Filipin staining, (G) *Salmonella* early interaction with and (H) invasion of K562 cells. Data is plotted as mean + sd of 6 (F, G) or 8 (H) replicates and statistical analysis was performed by Mann-Whitney U test (F) or 2-way ANOVA, whereby significance for the factor ‘treatment’ is indicated (G, H). (I-J) Simvastatin treatment for 48 h reduces (I) cholesterol levels and (J) *Salmonella* invasion. Data is plotted as mean + range or 3 replicates (I) or mean + sd of 9 replicates (J). Statistical analysis was performed by Mann-Whitney U test (I) or 2-way ANOVA, whereby significance for the factor ‘treatment’ is indicated (J). Dotted lines in C, F and I depict the MFI for unstained samples. *, p < 0.05; **, p < 0.01; ***, p < 0.001; MFI, median fluorescence intensity.

To determine the role of cholesterol levels independently of the cell cycle, methyl-β-cyclodextrin (MBCD) was used to extract cell surface cholesterol (57), which resulted in reduced Filipin staining (Fig 2F) with minimal effects on cell viability (Fig S2C-D). MBCD pretreatment of K562 reduced early interaction of *Salmonella* wt and Δ4 (Fig 2G), as well as invasion by *Salmonella* wt (Fig 2H). Moreover, we treated K562 cells with the statin drug Simvastatin for 48 h to block cholesterol biosynthesis (58), which did not affect cell viability (Fig S2E-F). Simvastatin treatment resulted in modestly reduced Filipin staining (Fig 2I) and, most notably, reduced *Salmonella* invasion (Fig 2J). Thus, while our results support that membrane cholesterol promotes *Salmonella* - host cell interaction (25, 30, 56, 59–61), suspension-grown mitotic cells appear not to be preferentially targeted by *Salmonella*. Rather, cholesterol-enriched cells display increased bacterial association throughout the cell cycle.

### Characterization of K562 gene expression patterns and cell surface features

Cell membranes targeted by *Salmonella in vivo* are typically covered by a complex glycocalyx, which is especially true for the apical intestinal epithelial surface that sports an ∼1 µm thick coat of heavily glycosylated transmembrane mucins and other glycoproteins (62). We therefore next quantified the expression of membrane-attached (glyco)proteins through global transcriptome and proteomes for unperturbed K562. Typical high abundance transcripts and proteins were detected with high expression (Fig S3A-D), providing a quality control of the datasets. Furthermore, inflammatory caspases were not expressed in K562 cells as opposed to non-transformed IECs derived from differentiated human 2D enteroid-derived monolayers (Fig S3E-H; (33)). This offers increased robustness of the K562 experimental system, since *Salmonella* invasion-induced cell death will be minimal.

Glycosaminoglycans (GAGs) are the largest glycans produced by animal cells and thus represent a substantial portion of the glycocalyx and play an important role at host-pathogen interfaces (63). Therefore, we assessed the presence of heparan sulfate (HS) and chondroitin sulfate (CS) GAGs, first by determining the expression levels of core proteins anchoring these glycan structures (Fig 3A-D). Compared to human 2D enteroid-derived IEC monolayers, our data indicated overall low expression levels of HS and CS core proteins (Fig 3A-D). This is in agreement with a previous study (64) and points towards a rudimentary GAG layer on the K562 surface. To confirm these data, we directly assessed the glycan portion of these proteoglycan structures. GAGs comprise a short tetrasaccharide linker attached to the core protein, elongated by linear chains of repeated and heavily sulfated disaccharide units, whereby the identity of the disaccharide units determines the GAG family (63). We quantified the amount of HS and CS disaccharides on K562 and analyzed their sulfation patterns (Fig 3E-F). In agreement with the transcriptome and proteome data, and earlier literature (64), this analysis confirmed very low levels of HS, and low levels of also CS (65). Due to the very low abundance of HS, the sulfation pattern was only assessed for CS, revealing that 4S sulfation was the most common type (Fig 3F). Altogether, these results indicate that the K562 surface features low levels of HS and CS proteoglycans at baseline and hint towards only a rudimentary glycocalyx.

**Figure 3.**
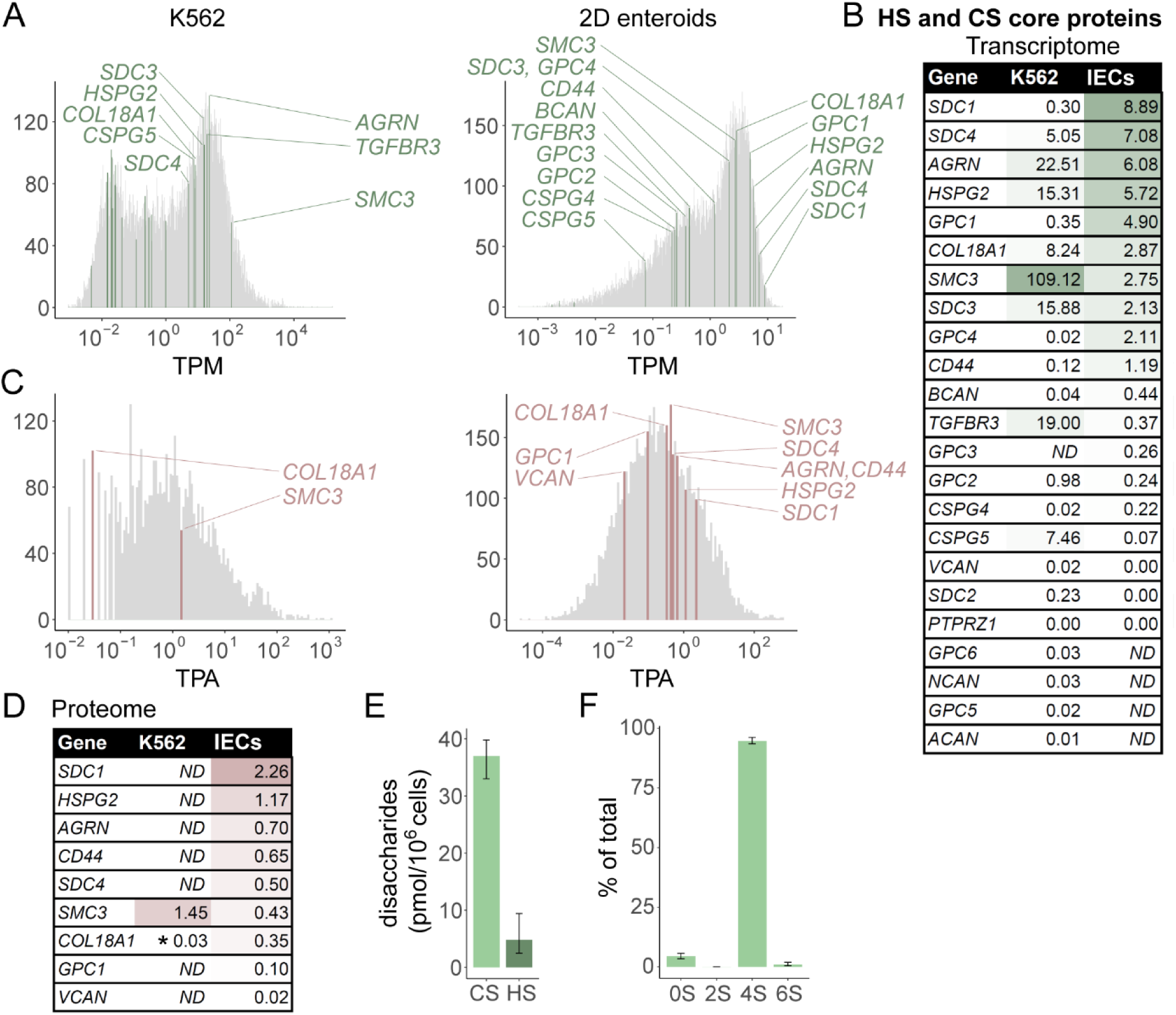
Characterization of K562 expression patterns and glycosaminoglycan content. (A-D) Transcriptome and proteome analysis of K562 cells at steady-state and comparison to differentiated human 2D enteroid-derived IEC monolayers (data from (33)). (A) Transcripts with TPM (transcript per kilobase million) values > 0 are plotted in a histogram, bins containing transcripts for HS or CS core proteins are highlighted. (B) List of TPM values for transcripts highlighted in A. The mean of 4 replicates is depicted for K562, while the mean of 3 replicates for independent human enteroid lines derived from 2 patients is shown for IECs. (C) Identified proteins (≥1 unique + razor peptide) with TPA (total protein amount in fmol/µg total protein) values > 0 are plotted in a histogram, bins containing HS or CS core proteins are highlighted. (D) List of proteins highlighted in C. The mean of 3 (IECs) or 4 (K562) replicates is depicted. Asterisks indicate identified proteins with <3 unique + razor peptides. (E-F) Analysis of (E) HS and CS disaccharides and (F) CS sulfation patterns plotted as mean + range of 6 replicates with 1-4x 10^7^ cells per replicate. ND, not detected.

### Host glycans and bacterial cell surface appendages obstruct *Salmonella* access to the host cell membrane

Building on this K562 surface characterization, we sought to determine the function of glycocalyx components as either barrier structures that hamper bacterial targeting, or as receptors for *Salmonella* adhesins promoting membrane access (10, 14–17, 20, 33, 40). Neuraminidase treatment was first used to trim glycan structures present on the K562 surface at steady state (Fig 4A) without affecting cell viability (Fig S2G-H). This promoted *Salmonella* adhesion to (Δ4; Fig 4B), early interaction with (wt; Fig 4B), and invasion of (Fig 4C) K562 cells. This suggests that even a rudimentary glycocalyx may represent a barrier for *Salmonella* host cell membrane targeting.

**Figure 4.**
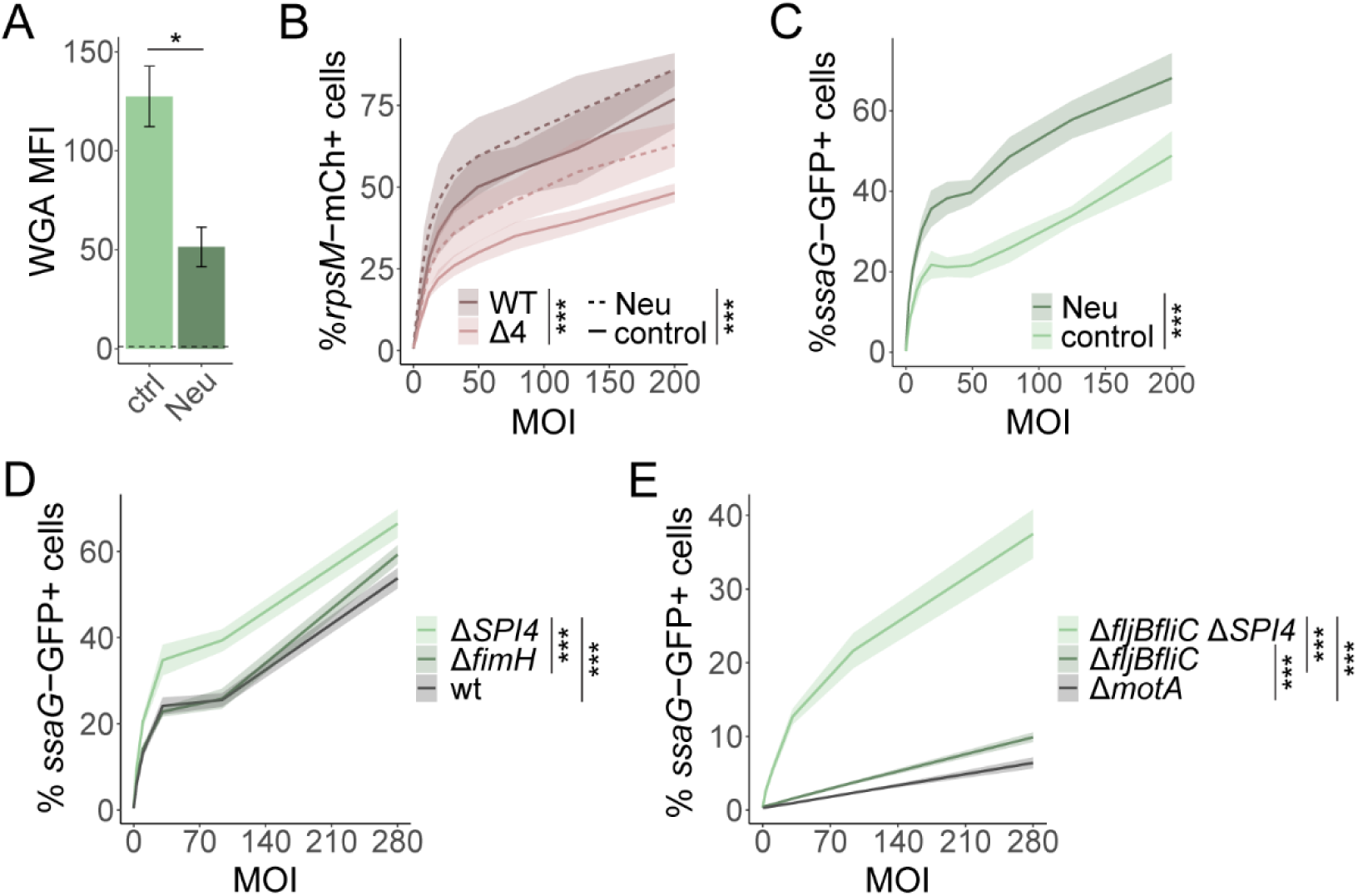
Host glycans and bacterial surface appendages obstruct *Salmonella* access to the host cell membrane. (A-B) Neuraminidase (Neu) treatment of K562 cells for 1 h (A) reduces cell surface glycosylation as assessed by Wheat germ agglutinin (WGA) staining, and increases (B) *Salmonella* wt and Δ4 early interaction and (C) *Salmonella* wt invasion of host cells. Data is plotted as mean + sd of 4 replicates (A), ≥10 replicates (B), or 8 replicates (C). Statistical analysis was performed by Mann-Whitney U test (A) or 2-way ANOVA (B-C), whereby significance for the factors ‘strain’ and ‘treatment’ is indicated. (D) Deletion of the large SiiE adhesin encoded on SPI4, but not removal of the fimbrial tip protein FimH, increases *Salmonella* invasion of K562 cells. (E) Deletion of flagellar subunits (Δ*fljBfliC*) increases *Salmonella* invasion efficiency in K562 cells compared to a non-motile, but still flagellated strain (Δ*motA*). Additional removal of SiiE (Δ*fljBfliC* Δ*SPI4*) further facilitates invasion. Infection was allowed to proceed for 60 min to allow non-motile bacterial strains to reach the K562 cells in the absence of centrifugation. In D-E, data is plotted as mean + sd of ≥ 5 replicates and statistical analysis was performed by 2-way ANOVA and TukeyHSD. *, p < 0.05; ***, p < 0.001; MFI, median fluorescence intensity.

Glycan moieties such as sialic acid (targeted by Neuraminidase) and mannose are, when presented on the right surface proteins, known binding sites for the *Salmonella* giant adhesin SiiE encoded on SPI4 (17) and the fimbrial tip protein FimH (66), respectively. Based on the observed barrier effect of the glycocalyx during *Salmonella* targeting of K562 cells, we next determined the role of glycan-binding adhesins SiiE and FimH in this context. In striking contrast to targeting of non-transformed human IECs (4), we found that SiiE actually impeded *Salmonella* invasion of K562 cells (Fig 4D). Moreover, the absence of FimH, which constitutes the tip of fimbrial adhesins, did not notably affect *Salmonella* invasion (Fig 4D). This led us to speculate that bulky bacterial cell surface structures such as SiiE might actually obstruct *Salmonella* access to the host cell membrane in the absence of optimal host cell surface target presentation. To test if this concept held true more broadly, we investigated the role of other bacterial surface appendages, the flagella, during K562 targeting. Deletion of flagellins (Δ*fljBfliC*) indeed resulted in improved invasion efficiency compared to non-motile, but still flagellated *Salmonella* (Δ*motA*; Fig 4E). Intriguingly, additional removal of SiiE (Δ*fljBfliC* Δ*SPI4*) further promoted invasion (Fig 4E), which is again in stark contrast to non-transformed human IECs, in which this strain is virtually non-invasive (4). This supports the notion that both host cell glycans and bacterial cell surface appendages obstruct access to the host cell membrane for T3SS-1-mediated invasion in the absence of cognate receptors for *Salmonella* adhesins. The presence or absence of specific glycoprotein structures in K562 cells, however, remained to be formally assessed.

### Even glycoproteins implicated as *Salmonella* adhesin receptors form steric barriers to membrane targeting

Non-transformed human epithelial cell subsets often express multiple transmembrane mucins, of which at least MUC1 and MUC13 can be targeted by *Salmonella* SiiE (20, 40). By global transcriptome and proteome profiling, such transmembrane mucins, as well as the M cell-associated GP2 glycoprotein (FimH target; (10)), were found expressed at or below the detection limit in K562 cells (Fig 5A-D). This substantiated our notions above, and suggested that ectopic expression of individual mucins/glycoproteins in K562 would allow to study their effect on *Salmonella* surface targeting in isolation.

**Figure 5.**
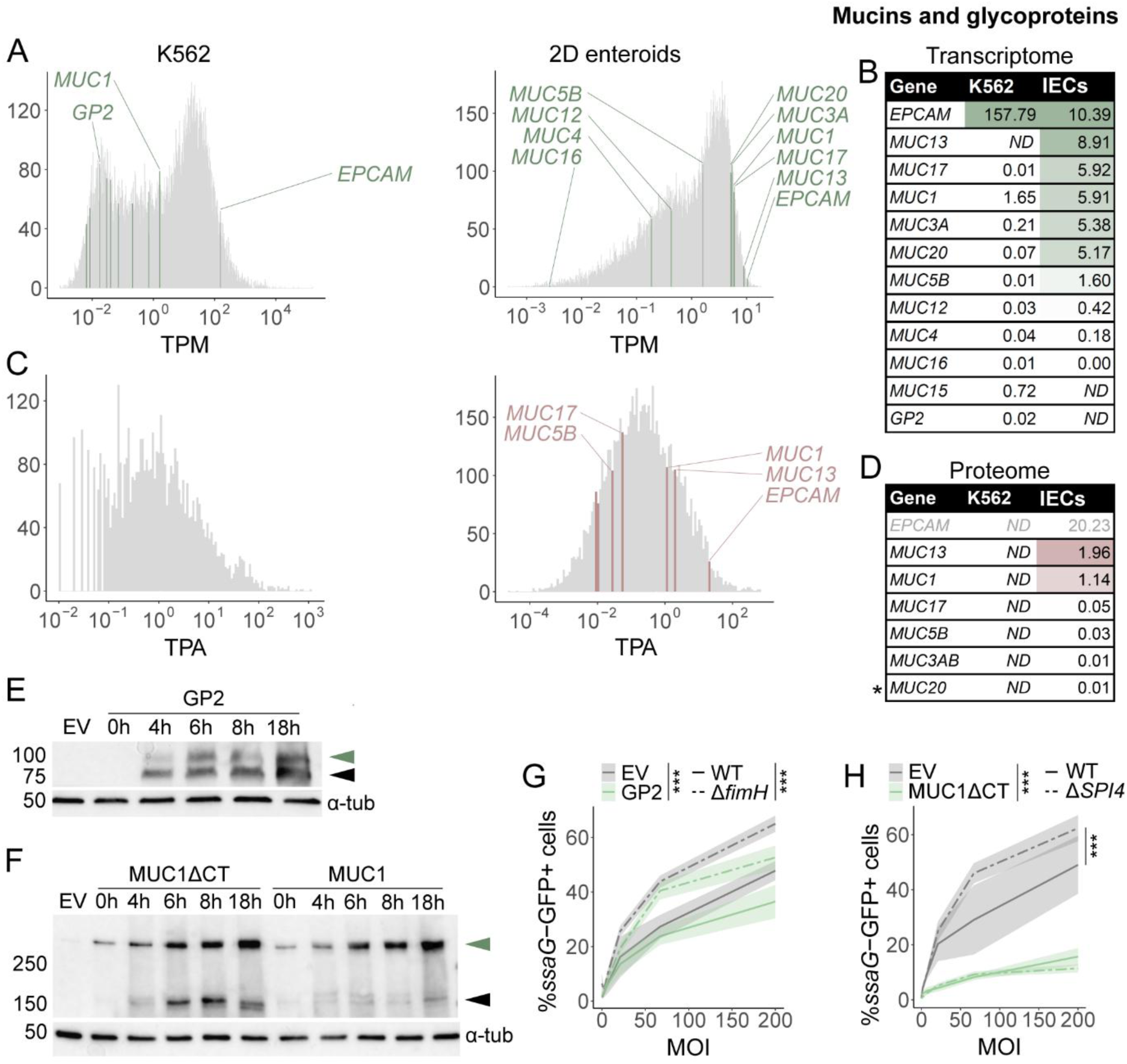
Cell surface glycoproteins form a steric barrier for *Salmonella* targeting. (A-D) Transcriptome (A-B) and proteome (C-D) analysis reveals that differentiated human IECs in 2D enteroid-derived monolayers express various transmembrane mucins and the epithelial marker EPCAM (data from (33)), whereas K562 cells display low to undetectable levels of mucin expression. (A) Transcripts with TPM > 0 are plotted in a histogram and bins containing transcripts for transmembrane mucins and other cell surface proteins are highlighted. (B) List of TPM values for transcripts highlighted in A. The mean of 4 replicates is depicted for K562, while the mean of 3 replicates for independent human enteroid lines derived from 2 patients is shown for IECs. (C) Identified proteins (≥1 unique + razor peptide) with TPA > 0 are plotted in a histogram, bins containing transmembrane mucins and other surface proteins are highlighted. (D) List of proteins highlighted in C. The mean of 3 (IECs) or 4 (K562) replicates is depicted. Asterisks indicate identified proteins with <3 unique + razor peptides. (E-F) Western blot analysis indicates successful ectopic expression and time-dependent glycosylation of GP2, MUC1 and MUC1ΔCT (lacking the cytoplasmic tail) in K562 cells. The expected sizes for non-glycosylated proteins, marked by black arrowheads, are ∼64 kDa for GP2 and ∼175 kDa for MUC1, whereas the glycosylated versions, marked by green arrowheads, are estimated to be ∼70-80kDa for GP2 and >250 kDa for MUC1. Time is indicated as hours post induction. (G) Ectopic expression of GP2 reduces invasion of *Salmonella* wt and Δ*fimH*. Data is plotted as mean + sd of 8 (wt) or 4 replicates (Δ*fimH*), whereby one replicate corresponds to an individually transfected culture infected with 2 different inocula (mean for bacterial replicates). (H) Ectopic expression the larger surface glycoprotein MUC1ΔCT strongly reduces *Salmonella* invasion, indicating that the glycocalyx forms a size- dependent barrier for host cell targeting. Data is plotted as mean + sd of 12 (wt) or 4 replicates (Δ*SPI4*), with replicates defined as in G. Statistical analysis was performed by 2-way ANOVA (G-H) and TukeyHSD (H). ***, p < 0.001; EV, empty vector.

Using the pMEP4 vector system (44–46), GP2 as well as full-length MUC1 and a version lacking the cytoplasmic tail (MUC1ΔCT) were successfully expressed and glycosylated in a time-dependent manner in K562 cells (Fig 5E-F, S4A-E). Of note, GP2 expression moderately reduced invasion of both *Salmonella* wt and Δ*fimH* (Fig 5G), illustrating that elevated GP2 if anything introduces a barrier for *Salmonella* K562 cell targeting despite its function as an adhesin target structure. Similar to GP2, but with dramatically larger effect size, MUC1 expression in K562 cells also reduced invasion by both wt and SiiE-deficient (Δ*SPI4*) *Salmonella* (Fig 5H, Fig S4F), again despite it being demonstrated as an interaction partner for SiiE. Confirming our previous data (Fig 4D-E), SiiE conferred a disadvantage for the invasion of K562 cells in the absence of MUC1, while invasion was low for both wt and Δ*SPI4 Salmonella* in MUC1-overexpressing cells (Fig 5H). Importantly, the more pronounced reduction in invasion in MUC1-compared to GP2-expressing cells (Fig 5G-H) is in line with the larger size of MUC1 and supports the conclusion that the glycocalyx functions as a size-dependent steric barrier for *Salmonella* targeting of the host cell membrane.

### Lectin stainings and structural prediction reveal physiologically relevant glycosylation of ectopically expressed MUC1

As ectopically expressed glycoproteins might forfeit their adhesin receptor function due to aberrant glycosylation patterns, we next sought to assess mucin-type O-glycosylation patterns in K562 cells. Stainings with a panel of lectins specific for core 1 mucin-type O-glycans, namely Jacalin, and peanut agglutinin (PNA) (67), or terminal glycan motives, namely *Sambucus nigra* agglutinin I (SNA-I), *Ulex europaeus* agglutinin I (UEA-I) and wheat germ agglutinin (WGA) (67), revealed increased signal upon ectopic MUC1 expression for Jacalin, PNA and WGA (Fig S5A-E). In line with this, transcriptome and proteome data confirmed that glycosyltransferases required for addition of the first N-acetylgalactosamine (GalNAc; *GALNT* gene family) moiety to serine/threonine of the extracellular variable number of tandem repeats (VNTR) domain, as well as for elongation with galactose to form core 1 structures (*C1GALT1*) were expressed in both human IECs and K562 cells (Fig S5F-I). Notably, the lectin staining pattern with close to background (empty vector control) intensities for SNA-I and UEA-I, selective for motifs containing terminal sialic acid and fucose residues, respectively (67), and a prominent WGA signal, detecting N-acetylglucosamine (GlcNAc), and to a lesser extent sialic acid and GalNAc (67, 68), is reminiscent of immature human enteroid-derived IEC monolayers (33). This hints towards physiologically relevant, although not fully mature, mucin-type O-glycosylation in MUC1-transfected K562. To gain further insights into specific mucin-type O glycan structures, we took advantage of an online tool to predict glycan sequences based on the expression levels of glycosyltransferases (69) in K562 and the previously published human IEC transcriptome data (33). The predictions confirmed the presence of Jacalin, PNA and WGA binding motifs in K562, whereas SNA-I and UEA-I binding motifs were restricted to IECs (Fig S6). As expected, this analysis also pointed to higher structural diversity of mucin-type O-glycans in IECs (Fig S6). Structures predicted for K562 cells agreed with previous descriptions of this cell line by others (70) and, most notably, represented a subset of O-glycans in differentiated human IECs. Despite the absence of SNA-I binding motifs, most of the K562 glycans contained terminal sialic acid residues, which have been suggested to interact with the *Salmonella* SiiE adhesin (17, 20). Altogether, these data support physiologically relevant O-glycosylation of MUC1 ectopically expressed in K562 cells.

### The glycocalyx forms a size-dependent barrier towards *Salmonella* targeting of the host cell membrane

Flow cytometry analysis of MUC1ΔCT-transfected K562 cells stained with WGA indicated a subpopulation (∼40%) that did not express high levels of glycosylated MUC1 (Fig 6A-B). Strikingly, a *Salmonella* invasion assay combined with subsequent WGA staining revealed that virtually only this WGA^low^ subpopulation (with WGA intensity comparable to empty vector control transfectants) was efficiently targeted by *Salmonella* (Fig 6C-E). To assess whether the phenotype of MUC1-expressing cells can be reverted, we treated transfected K562 cells with the mucinase StcE, a protease that specifically cleaves off the extracellular, glycan-bearing domain of mucins (71). Indeed, StcE treatment reduced WGA staining to the level of the empty vector control (Fig 6F) without affecting cell viability (Fig S4G-H). Most importantly, mucinase treatment also restored *Salmonella* wt and *ΔSPI4* invasion of MUC1-expressing cells (Fig 6G).

**Figure 6.**
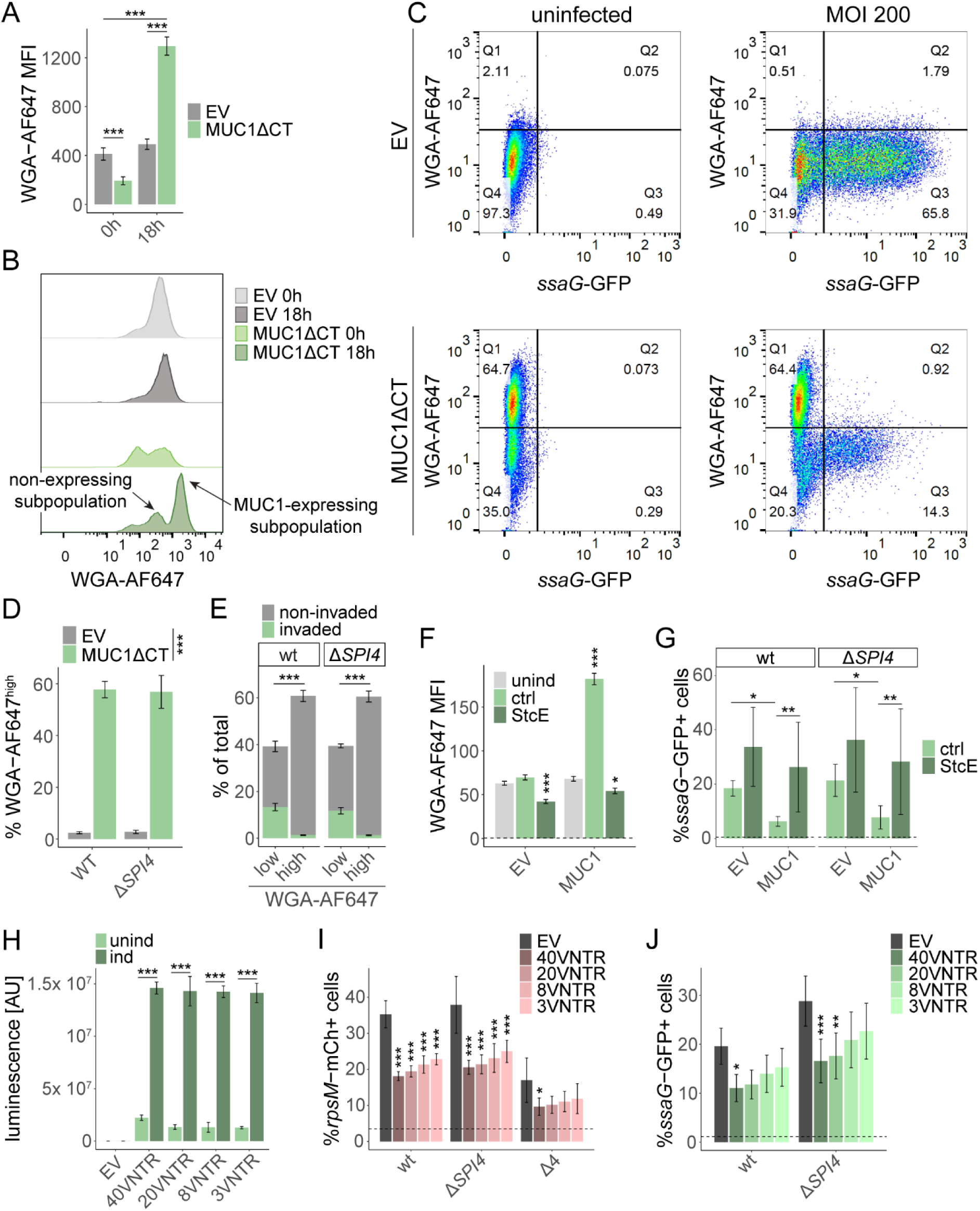
Ectopic MUC1 imposes a size-dependent constraint on *Salmonella* surface targeting. (A) WGA staining increases upon induced (18h) ectopic expression of MUC1. Data is plotted as mean + sd of 4 replicates. (B) A subpopulation of transfected cells does not successfully express MUC1 upon induction, as indicated by low WGA intensity comparable to background. (C-E) WGA staining following *Salmonella* infection of transfected and induced cells indicates that almost exclusively the non-expressing, WGA^low^ subpopulation is targeted in MUC1ΔCT-transfected cells. (C) Virtually no *ssaG*-GFP+ WGA^high^ double-positive cells (Q2) are observed in MUC1ΔCT-transfected cells infected with *Salmonella* wt. (D) WGA^high^ cells are specifically detected in MUC1ΔCT-transfected samples. Pooled data for all MOIs is depicted. (E) Quantification of population sizes illustrated in C confirms that cells successfully expressing MUC1 largely remain uninvaded. Data for MOI 200 is depicted. In D- E, data is plotted as mean + sd of 4 replicates, whereby one replicate corresponds to an individually transfected culture infected with 2 different inocula (mean for bacterial replicates). (F-G) Treatment of MUC1-transfected K562 cells with the mucinase StcE (F) reverts increased cell surface glycosylation as assessed by WGA staining, and (G) restores *Salmonella* invasion (MOI 50). Data is plotted as mean + sd of 4 (F) or 7 (G) replicates. (H-J) K562 cells ectopically expressing HiBiT-tagged MUC1 versions with varying numbers of VNTRs, resulting in different lengths, were infected with *Salmonella* at MOI 50. (H) Luminesence-based detection of HiBiT-tagged MUC1 versions reveals robust and comparable expression of all constructs upon induction. (I) Early interaction and (J) invasion assays illustrate the size-dependent barrier effect of MUC1 towards *Salmonella* targeting of the K562 host cell surface. Data is plotted as mean + sd of 4 (H) or 6 (I-J) replicates, whereby one replicate is defined as described above. Statistical analysis was performed by 2-way ANOVA (A, D-J) and TukeyHSD (A, E-J). *, p < 0.05; **, p < 0.01; ***, p < 0.001; MFI, median fluorescence intensity; EV, empty vector; AU, arbitrary units.

To further confirm the size-dependency of the glycocalyx barrier, we expressed MUC1 structures of different length on the surface of K562 cells by varying the number of VNTRs from 3 to 40, whereby the longest version corresponds to the previously used MUC1ΔCT (Fig S4I-K). These MUC1 versions were designed to contain a short N-terminal HiBiT tag (72) between the signal sequence cleavage site and the VNTR domain, which is predicted to not affect protein folding. Importantly, this allows for lectin-independent detection of ectopic protein expression via a luminescence assay, as glycosylation levels are dependent on the VNTR domain length (Fig S4K). Indeed, luminescence-based detection revealed robust and comparable expression of all MUC1 constructs upon induction (Fig 6H). Most notably, both the *Salmonella* early interaction assay and the invasion assay revealed a clear size-dependent suppressive effect of the number of MUC1ΔCT VNTRs on bacterial host cell surface targeting (Fig 6I-J). Altogether, these findings demonstrate that cell surface glycoproteins, including also potential receptors for *Salmonella* adhesins, form size-dependent obstacles towards *Salmonella* targeting when presented on a cell membrane with an otherwise minimal glycocalyx.

## DISCUSSION

While the development of physiology-mimicking organotypic and organ-on-a-chip technologies is revolutionizing the study of enteropathogenic infections (73–79), they build upon the knowledge generated by decades of continuous cell line culture-based research. This highlights such models as still highly valuable tools to study the mechanistic and molecular underpinnings of host-pathogen interactions, with far superior possibilities for genetic perturbation, bottom-up approaches, and large-scale screening. With the K562 model developed herein, we have expanded the repertoire of available infection models with a platform optimally suited for flow cytometry-based *Salmonella* binding and invasion assays, coupled with controllable ectopic expression of host factors potentially involved in host-pathogen interplay.

Employing this model to determine host cell characteristics affecting *Salmonella* targeting of the host cell surface, we revisited a long-standing debate about the mechanism underlying preferential invasion of mitotic cells and the links to concomitant changes in cholesterol levels and cell shape (2, 55, 56). Our data confirm increased cholesterol levels during mitosis and preferential *Salmonella* targeting of a cholesterol-rich host cell membrane (25, 30, 56, 59–61). However, we found that cholesterol-rich cells display increased association with *Salmonella* throughout the cell cycle. Cholesterol may have multiple roles during *Salmonella* targeting. It directly binds the T3SS-1 tip protein SipB to enhance T3SS-1 docking and promote effector translocation (3, 25, 26), but also affects physical and biological membrane properties and mediates the formation of specialized membrane domains (lipid rafts) which differ in their lipid composition and cluster certain transmembrane proteins of relevance for infection (80, 81). The contribution of cholesterol, and potentially its precursors, to host cell targeting is therefore likely a function of all these aspects, as the specific absence of cholesterol has been indicated to not affect *Salmonella* invasion when all its upstream precursors were preserved (82).

Cell surface glycostructures constitute the initial bacterial contact point with e.g. the intestinal epithelial surface and hence have been implied in various interactions with enterobacterial pathogens (10, 14–17, 20–22, 31–36, 39–43, 83–86). Thereby, glycans have either been implied as binding targets for bacterial adhesins (10, 14–17, 20–22, 40, 42, 43) or the glycocalyx has been assigned a barrier function towards bacterial pathogens and commensals (31–36, 38–41). Deglycosylation using Neuraminidase indicated that already the modest endogenous K562 glycocalyx forms a barrier towards membrane-attacking *Salmonella*. GAGs are important constituents of the glycocalyx and have been implicated as receptors for *Salmonella* and *E. coli* host cell targeting (16, 42), but appear minimally produced by K562 cells. Due to the multitude of core proteins and series of glycosyltransferases required for expression and glycosylation of these structures, ectopic expression of proteoglycans was not deemed feasible in our model.

Instead, we focused on specific glycoproteins and transmembrane mucins, which constitute key components of the intestinal glycocalyx (62, 87), and have been successfully expressed in other cell models (36, 88). Ectopic expression of GP2 and MUC1 in K562 cells notably reduced, rather than increased, *Salmonella* host cell targeting in a glycoprotein size-dependent manner. This suggests that when presented on an otherwise essentially bare surface, intestinal-relevant elongated glycoproteins such as MUC1 represent predominantly a barrier for host cell targeting. This is further supported by our data using treatment with the mucinase StcE, or shortened MUC1 versions, which both reduced this barrier effect. In fact, even bacterial appendages including the giant SiiE adhesin itself restrict interaction of T3SS-1-expressing *Salmonella* with K562 cells, which is in stark contrast to the important role for SiiE in docking to the apical surface of primary human IECs (4) with its dense glycocalyx. Our analysis of mucin-type O-glycosylation patterns indicated that MUC1 expressed in K562 cells is glycosylated in a physiologically relevant fashion, albeit with reduced structural diversity. A combination of lectin stainings and glycan sequence prediction further suggests that mucin-type O-glycosylation in K562 cells resembles that in immature, proliferative human IECs (33). Nevertheless, the predicted presence of terminal sialic acid moieties, which are the reported SiiE binding motif (17), favors the presence of receptor-competent MUC1 in transfected K562 cells. However, the fact that the barrier effect was similarly observed for both *Salmonella* wt and mutants lacking GP2- and MUC1-binding adhesins (Δ*fimH* and Δ*SPI4*, respectively), implies that adhesin-receptor interactions do not further promote targeting in the context of an essentially bare membrane with a low-complexity glycocalyx.

These data support a central role of transmembrane mucins as a barrier towards pathogenic (and commensal) bacteria as described in some Caco-2 cell studies (36), human enteroid/colonoid-derived monolayers (33) and the murine intestine *in vivo* (32, 38, 40, 41). The glycocalyx barrier formed by transmembrane mucins has been shown to be fortified during IEC differentiation (33) and during weaning in mice (38). As IECs differentiate along the crypt-villus axis, bacterial encounter increases, as it does during suckling-to-weaning transition with the enhanced uptake of environmental microbes. This spatial and temporal fortification of the glycocalyx upon increased frequency of bacterial encounter further highlights the central role of transmembrane mucins as a steric barrier during host cell targeting by various pathogens.

Altogether, our results establish a poorly glycosylated, bare membrane rich in cholesterol and with minimal presence of elongated glycoproteins as the optimal targeting surface for virulence-primed *Salmonella* and suggest that the requirement for adhesin-receptor interactions when targeting such a simple surface is minimal. Nevertheless, adhesins are important virulence factors (4, 20, 27, 34), and likely the result of an ongoing arms race as enterobacterial pathogens are faced with a largely impenetrable glycocalyx at e.g. the intestinal epithelial surface. In turn, IECs have been shown to release cleaved transmembrane mucins as decoy receptors to counter bacterial onslaught (34, 40), and enterobacterial pathogens even employ various adhesin-independent strategies to overcome the glycocalyx barrier (31, 39, 75, 77, 89–91). We anticipate that the model developed herein will be a valuable tool to further investigate the intricacies of the glycocalyx barrier(s) and mechanistically characterize barrier phenotypes observed in more complex organotypic or *in vivo* models. The high-throughput K562 model can for example enable combined expression of multiple cell surface glycoproteins at ratios mirroring the composition across the different organs and cell types targeted by *Salmonella* (or other microbes) during *in vivo* infection.

## MATERIALS AND METHODS

### Bacterial strains, plasmids and culture conditions

Bacterial strains and reporter plasmids used in this study are listed in Table S1. All *Salmonella enter*ica serovar Typhimurium (*Salmonella*, *S*.Tm) strains were of SL1344 background (SB300, Streptomycin resistant, Sm^R^) (92). The wildtype (wt) and previously described mutants Δ*invG*, Δ*sipA*, Δ*sopBEE2*, Δ*sipA* Δ*sopBEE2* (Δ4), Δ*SPI4*, Δ*motA*, Δ*fljBfliC* and Δ*fljBfliC* Δ*SPI4* were used. The Δ*fimH* mutant was generated by transfer of a previously described deletion from *Salmonella* 14028 (C0831) (93) by P22 transduction. The genotype was verified by PCR using the primer pairs k1/fimH scr F, k2/fimH scr R and fimH scr F/fimH scr R (Table S2). The intracellular vacuolar, SPI2-dependent pM975 (p*ssaG*-GFPmut2) (47, 48), cytosolic pZ1400 (p*uhpT*-GFP) (4, 49) and constitutive pFPV-mCherry (p*rpsM*-mCherry; Addgene plasmid number 20956) (52) reporter plasmids (Table S1) have been described and validated previously. For K562 and HeLa infections, *Salmonella* was grown overnight at 37°C for 12 h in LB/0.3 M NaCl (Sigma-Aldrich) with appropriate antibiotics and sub-cultured (1:20 dilution) in the same medium without antibiotics at 37°C for 4 h. The inoculum was reconstituted in complete RPMI or infection medium (Table S3) for K562 cell infections, or complete DMEM (Table S3) for HeLa cell infections. Where required, reconstituted inocula were further serially diluted. *Escherichia coli* Dh10b used for amplification of pMEP4 vectors and the CMV-EBNA1 plasmid was cultured in LB supplemented with appropriate antibiotics.

### K562 and HeLa cell culture and cell surface manipulations

K562 CCL-243 cells (ATCC) were maintained in complete RPMI (Table S3) supplemented with 1x PenStrep (Gibco) at 37°C, 5% CO_2_ in slightly tilted cell culture flasks at densities between 50’000 to 500’000 cells/mL. Cultures were passaged every 2-3 days when the density exceeded 500’000 cells/mL, at a ratio ranging from 1:4-1:12. Cultures from passages 7-54 were used for experimentation. HeLa CCL-2 cells (ATCC) were cultured in complete DMEM (Table S3) supplemented with 1x PenStrep at 37°C, 10% CO_2_ and passaged every 2-3 days when the confluency reached 70% or more. Cells were washed once with DPBS (Gibco), detached using Trypsin-EDTA (Gibco) and reconstituted in fresh medium at a dilution ratio between 1:2-1:8 for seeding in new cell culture flasks. Passages 4-30 were used for experimentation. Where indicated, cells were treated with 0.5 µg/mL Taxol (SIGMA-Aldrich) in complete RPMI for ∼18 h (starting with 300’000 cells/mL), 2 mM methyl-β-cyclodextrin (MBCD; SIGMA-Aldrich) in infection medium for 30 min (10^6^ cells/mL), 5 µM Simvastatin (SIGMA-Aldrich) in complete RPMI for 48 h (starting with 250’000 cells/mL) or 0.1 U/mL Neuraminidase (SIGMA-Aldrich) in complete RPMI for 1 h (∼500’000 cells/mL).

### Ectopic protein expression via K562 transfection with pMEP4

pMEP4 vectors (44–46) with the inserted sequences for expression of GP2, MUC1 and a truncated version of MUC1 lacking the cytoplasmic tail (MUC1ΔCT) were ordered from GenScript using their subcloning service. The insert sequences designed for cloning into the multiple cloning site of pMEP4 via the restriction enzymes HindIII and BamHI are listed in Table S4. Sequences for MUC1 versions with varying number of VNTRs were designed to include a short N-terminal HiBiT tag, VSGWRLFKKIS, (72) between the signal sequence cleavage site and the VNTR domain as highlighted in Table S4. Upon delivery, pMEP4 vectors were reconstituted at 200 ng/μL in nuclease-free water (Thermo Fisher) according to GenScript’s instructions. 1 μL of the resuspended vector was used for heat shock transformation into chemically competent (0.1 M calcium chloride) *E. coli* Dh10b. Large amounts of pMEP4 and CMV-EBNA1 plasmids at high concentrations were obtained using GeneJET Plasmid Maxiprep kit (Thermo Fisher) according to the manufacturer’s protocol. For transfections (informed by (44–46)), K562 cells were reconstituted in room temperature (RT) electroporation medium (Table S3) at a concentration of 4x 10^7^ cells/mL. 2x 10^7^ cells were used for transfection via addition of transfection mix (8 µg of the respective pMEP4 vector, 5 µg CMV-EBNA1, 0.15 M NaCl), transfer to 0.4 cm cuvettes (BioRad) and electroporation at 340 V, 0.95 mF using a Gene Pulser II (BioRad). After 5 min, transfected cells were transferred to Leighton tubes containing 8 mL revitalization medium (Table S3) and incubated overnight at 37°C, 5% CO2. Selection was initiated by transfer of transfected cells to 30 mL selection medium (Table S3). After one week, cells were washed once in revitalization medium and ectopic expression was induced by transfer to induction medium (Table S3) at a concentration of 10^6^ cells/mL and incubation for ∼18 h unless otherwise indicated.

### Salmonella infections

For infection, K562 cells were reconstituted in complete RPMI or infection medium at a density of 5.56x 10^5^ cells/mL, and 180 µL were transferred to the wells of a round-bottom 96-well plate. Adjusted volumes and concentrations were used for infections in 15 mL tubes (4 mL, 250’000 cells/mL), FACS tubes (2 mL, 500’000 cells/mL), 6-well (2 mL, 375’000 cells/mL) or 12-well plates (2 mL, 500’000 cells/mL). Where applicable, Taxol was added back to the prepared cells prior to infection. HeLa cells were seeded at a density of 250’000 cells/well in complete DMEM in 6-well plates and incubated overnight for infection the day after. The *Salmonella* inoculum was prepared as described above and added to the cells at the indicated multiplicity of infection (MOI) to initiate infection. For early interaction assays, bacteria and host cells were co-incubated for 15 min unless otherwise indicated. For invasion assays, bacteria and host cells were co-incubated for 30 min unless otherwise indicated, followed by centrifugation at 100 g for 1 min and reconstitution in complete RPMI for K562 or complete DMEM for HeLa, respectively, containing 100 µg/mL Gentamicin and incubation for another 3-3.5 h. For 18 h infections, the Gentamicin concentration was reduced to 25 µg/mL at 2 h p.i. At 15 min or 3.5-4 h p.i., respectively, K562 cells were washed once (invasion assay) or twice (early interaction assay) in DPBS/1% BSA using gentle centrifugation at 100 g for 1 min to efficiently eliminate non-adherent bacteria. For infections in 6- or 12-well plates, K562 cells were transferred to FACS tubes, and HeLa cells were detached using Trypsin-EDTA, reconstituted in complete DMEM and transferred to FACS tubes for the wash steps. Following fixation with DPBS/2% paraformaldehyde (PFA) for 20 min at RT in the dark, the cells were washed one more time as described above and kept at 4°C until further analysis.

### Microscopy

Images were acquired on either an inverted Ti-eclipse microscope (Nikon) with a 60×/1.4 NA PLAN APO objective (0.19mm working distance; Nikon) and an Andor Zyla sCMOS camera (Abingdon, Oxfordshire, UK) with pixel size of 108 nm (for Fig 1A), or on a custom-built microscope based on an Eclipse Ti2 body (Nikon), equipped with 60x/0.7 and 40x/0.6 Plan Apo Lambda air objectives (Nikon) and a back-lit sCMOS (scientific complementary metal oxide semiconductor) camera with pixel size 11 µm (Prime 95B; Photometrics). Bright-field image acquisition was performed by differential interference contrast (DIC), and fluorescence imaging by the excitation light engine Spectra-X (Lumencor) and emission collection through a quadruple band pass filter (89402; Chroma). Fiji, a version of ImageJ (94) was applied for image analysis.

### Staining and flow cytometry analysis

For live-dead staining, cells were reconstituted in DPBS/2 µg/mL propidium iodide (PI; SIGMA-Aldrich) and kept on ice until flow cytometry analysis. 0.1% Triton-X-100 (Tx-100; SIGMA-Aldrich) was added to positive controls. For analysis of cell cycle phases, cells were reconstituted in DPBS/0.1% Tx-100/10 µg/mL PI/10 µg/mL RNase A (Thermo Fisher), incubated for 20 min at RT in the dark and kept at 4°C until flow cytometry analysis within 24 h. For Filipin staining, cells were fixed in PBS/4% PFA for 20 min at RT in the dark, washed once in DPBS/0.1% BSA and reconstituted in DPBS/0.1% BSA/80 µg/mL Filipin (SIGMA-Aldrich, F9765). Following incubation for 20 min at RT in the dark, cells were washed twice as described above and kept at 4°C until flow cytometry analysis. For lectin stainings, cells were fixed in PBS/2% PFA for 20 min in the dark, washed once in DPBS/0.1% BSA and reconstituted in DPBS/0.1% BSA containing the respective lectin at the dilution indicated in Table S5. Following incubation for 20 min at RT in the dark, cells were washed twice as described above and kept at 4°C until flow cytometry analysis. For antibody staining, cells were washed once in DPBS/0.1% BSA, fixed in DPBS/2% PFA for 20 min at RT in the dark, blocked with PBS/3% BSA for 30 min shaking and incubated with the respective primary antibody (Table S5) diluted in DPBS/3% BSA for 40 min at RT, shaking in the dark. After two additional washes in DPBS/0.1% BSA, samples were incubated with the respective secondary antibody (Table S5) diluted in DPBS/0.1% BSA for 40 min at RT, shaking in the dark. Following two additional washes with DPBS/0.1% BSA, samples were kept at 4°C until flow cytometry analysis. Flow cytometry was performed using a MACSQuant VYB (Miltenyi Biotec) and data was analyzed in FlowJo version 10.10.0 (95). Gating for single cells was performed based on an FSC-H versus FSC-A plot.

### Glycosaminoglycan analysis

Glycosaminoglycans were isolated from K562 cells as previously described (96) and subjected to digestion with chondroitinase ABC and heparinase I, II and III to generate disaccharides. Disaccharide composition and amounts were determined by RPIP-HPLC separation followed by postcolumn derivatization with cyanoacetamide and quantification in a fluorescence detector (97).

### Western blot

Cells were washed once in DPBS and pellets were stored at -80°C until further analysis. Frozen pellets were reconstituted in cold RIPA lysis buffer (Thermo Fisher) supplemented with cOmplete^TM^ EDTA-free protease inhibitor (Roche) at a concentration of 10^7^ cells/mL. After lysis for 10 min on ice, cells were centrifuged at 15’000 g, 4°C for 15 min to remove cell debris and supernatants were transferred to fresh tubes. A 1:10 dilution of the samples was used to determine protein concentrations in a bicinchoninic acid assay (Thermo Fisher) using dilutions of Pierce^TM^ BSA standard (Thermo Fisher) for reference. 20 µg of protein were mixed with 4x Laemmli buffer (Biorad) containing 5% β-mercaptoethanol, incubated for 5 min at 95°C and loaded on a 4–20% Mini-PROTEAN® TGX Stain-Free™ Protein Gel (Biorad) along with 5 µL of ladder (Biorad). Gels were run for 35 min at 200 V and transferred to a nitrocellulose membrane (0.2 μm pore size; Biorad) with a Trans-Blot Turbo Transfer System (Biorad) using the program for high molecular weight. A Gel Doc EZ Imager (Biorad) was used for 1 min activation of the gel and imaging of total protein on the blot. Blots were blocked in DBPS/0.1% Tween-20 (SIGMA-Aldrich)/5% milk for 1 h shaking at RT and incubated with the respective primary antibody (Table S5) diluted in DPBS/0.1% Tween-20/2% milk overnight shaking at 4°C. Thereafter, blots were washed three times in DPBS/0.1% Tween-20 for 10 min shaking at RT and incubated with the respective secondary antibody (Table S5) diluted in in DPBS/0.1% Tween-20 for 1 h shaking at RT. The membranes were washed another 3 times as described above, incubated with ECL Prime Western Blotting Reagent (Cytiva) for 5 min shaking at RT in the dark and imaged using a ChemiDoc MP (Biorad).

### HiBiT extracellular detection

Extracellular detection of HiBiT-tagged MUC1 versions at the surface of transfected K562 cells was conducted using the Nano-Glo® HiBiT Extracellular Detection System (Promega, N2420) according to the manufacturer’s instructions. In short, 10^4^ cells were diluted into a total volume of 50 µL RPMI without FBS in a white flat-bottom 96-well plate. After equilibration to RT, an equal volume of the reconstituted HiBiT extracellular detection reagent was added to each well. Luminescence was acquired 10 min after addition of the detection reagent using a TECAN SPARK microplate multimode reader.

### Transcriptome analysis

Library preparation and RNA sequencing was performed by the SNP&SEQ Technology Platform in Uppsala. Libraries were prepared from 1 μg total RNA using the TruSeq stranded total RNA library preparation kit with RiboZero Gold treatment (Illumina Inc. 20020598/9). The library preparation was performed according to the manufacturers’ protocol (#1000000040499). Analysis of RNA-seq data was performed using the best practice pipeline nf-core/rnaseq (https://github.com/nf-core/rnaseq). The workflow processes raw data from FastQ files, aligns the reads, generates gene counts, and performs extensive quality-control on the results. Detailed information about analysis pipeline can be found here: https://github.com/nf-core/rnaseq/blob/master/docs/output.md. Transcriptome data from differentiated human enteroid-derived IEC monolayers cultured in in-house made ENR differentiation medium (33) were used for comparison.

### Global proteomics of K562 cells

K562 cells were cultured as described. Four replicates (10^7^ cells grown for 2 days) were used for global proteomics analysis. Cell pellets were collected and lysed in a lysis buffer (50 mM DTT, 2 %(w/v) SDS, 100 mM Tris/HCl pH 7.8). After incubation at 95 °C for 5 min, the lysates were sonicated with 20 pulses of 1 s, 20% amplitude by using a sonicator coupled with a microtip probe. For digestion of proteins to peptides, MED-FASP method was performed using trypsin and Lys-C (98). C18 stage tips were used for desalting the peptide mixture and samples were stored at −20°C until analysis. A tryptophan fluorescence assay was performed to determine protein and peptide content (99). The global proteomics analysis was performed on a Q Exactive HF mass spectrometer (MS) (Thermo Fisher Scientific) coupled to a nano–liquid chromatography (nLC). An EASY-spray C18-column (50 cm long, 75 μm inner diameter) was used to separate peptides on an acetonitrile/water gradient with 0.1% formic acid over 150 min. MS was set to data dependent acquisition with a Top-N method (full MS followed by ddMS2 scans). MaxQuant software (version 2.1.0.0) with the human proteome reference from UniProtKB (October, 2022) were used to identify proteins. Total protein approach was used for protein quantification (98). Proteomics data generated in this study have been deposited in the PRIDE database and will be made public upon publication. Proteome data from differentiated human enteroid-derived IEC monolayers cultured in commercially available cENR (IntestiCult™ Organoid Differentiation Medium (Human), StemCell) (33) (PRIDE accession code PXD063020) were used for comparison.

### Statistical analysis

Data was analyzed and plotted using the tidyverse collection of R packages (100) in RStudio (101). Data is always plotted as mean + range for ≤3 replicates, and as mean + sd for ≥4 replicates. The statistical tests used are specified in the respective figure legend.

## ACKNOWLEDGEMENTS

We are grateful to members of the Sellin laboratory for helpful discussions. We thank Martin Gullberg for advice on pMEP vector designs, Inger Eriksson for conducting the HS/CS analysis, and Petra Muir, Susan Schlegel and Hanna Eriksson for advice regarding and maintenance of flow cytometry instrumentation. This work was supported by the Swedish research council (grants 2018-02223 and 2022-01590 to M.E.S., and grant 2020-01586 to P.A.), the Swedish Foundation for Strategic Research (grant FFL18-0165), and the SciLifeLab Fellows program to M.E.S. M.C. acknowledges a PhD scholarship from the Ministry of National Education of the Turkish Republic. Transcriptome analysis was done at the SNP&SEQ Technology Platform in Uppsala, part of the National Genomics Infrastructure (NGI) Sweden and Science for Life Laboratory. The SNP&SEQ Platform is also supported by the Swedish Research Council and the Knut and Alice Wallenberg Foundation.

## DISCLOSURE OF INTEREST

The authors declare no competing interests.

## AUTHOR CONTRIBUTIONS

Conceptualization: P.G., M.E.S.; methodology: P.G., J.W., M.C., W.B., C.v.B., L.K.; investigation: P.G., J.W., M.C., W.B., L.K.; formal analysis: P.G., J.W., M.C., W.B., C.v.B.; interpretation: P.G., J.W., M.C., P.A., T.P., L.K., M.E.S.; resources: T.P.; supervision: M.E.S., PA.; project administration: P.G., M.E.S.; funding acquisition: M.E.S., visualization: P.G., writing – original draft: P.G., M.E.S.; writing – reviewing and editing: all authors.

---

**Supplementary Figure S1.**
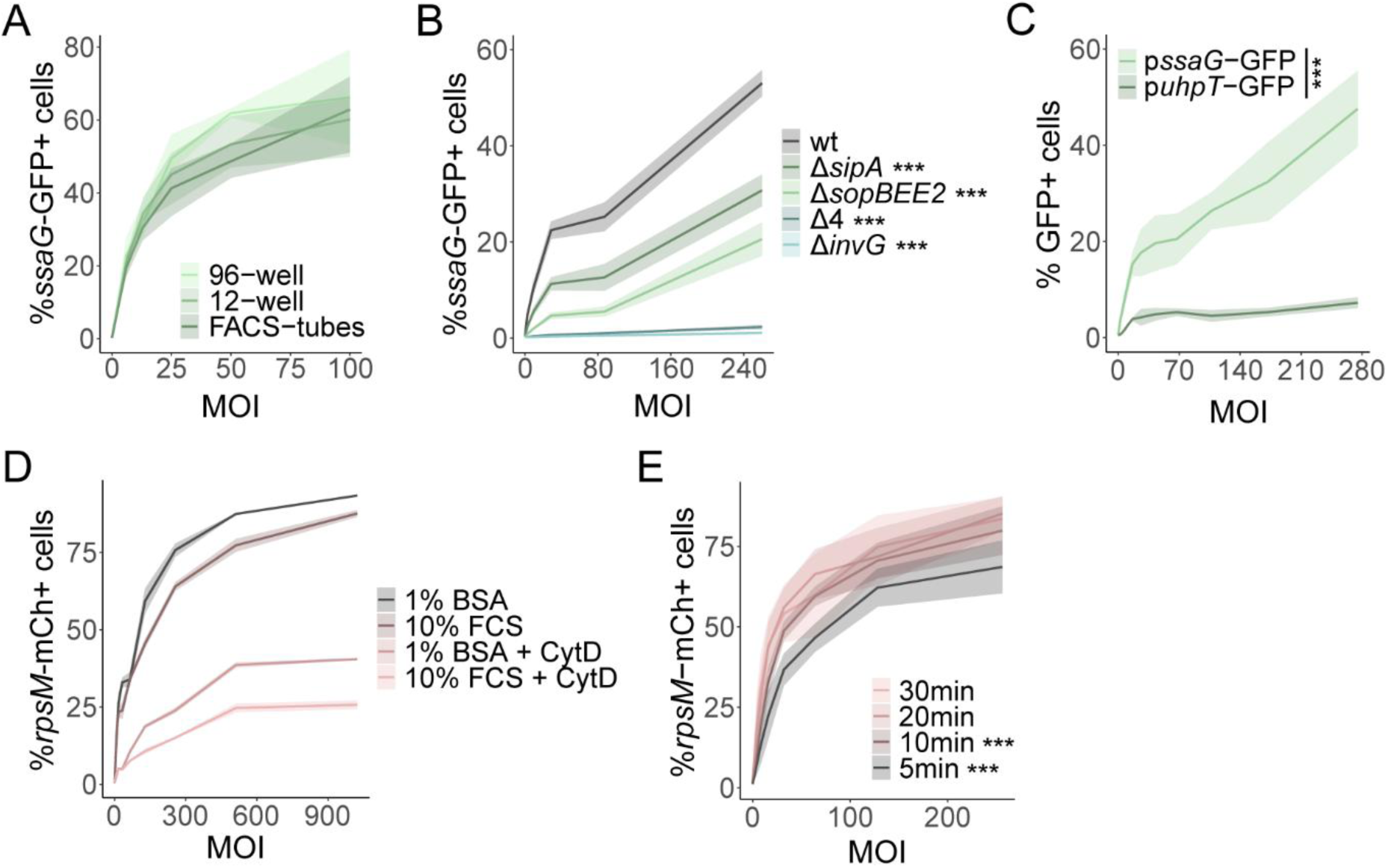
Validation of the high-throughput *Salmonella* infection model in K562 cells. (A) K562 infection with *Salmonella* (*S*.Tm) wt p*ssaG*-GFP for 30 min in different formats yields comparable invasion efficiencies. (B) Infection in 96-well plates allows for a high throughput invasion assay, in which multiple strains and MOIs can be compared in a single experiment. (C) An invasion assay with the vacuolar p*ssaG*-GFP and cytosolic p*uhpT*-GFP reporters reveals a small cytosolic *Salmonella* subpopulation in K562 cells. (D) An adhesion assay using Cytochalasin-D (CytD) treatment to block uptake of *Salmonella* wt results in low numbers of cell-associated bacteria regardless of the media type. Different media were tested because the presence of serum might reduce bacterial association with host cells, and hence supplementation with BSA is preferable. 10% FCS: complete RPMI (RPMI/10% FBS/1 mM NaPyr; see Table S3); 1% BSA: infection medium (RMPI/1% BSA; see Table S3). (E) Effect of infection time on *Salmonella* association with K562 cells to establish an early interaction assay. Data is plotted as mean + range of 2 (D) or 3 (A) replicates, or as mean + sd of 6-10 replicates (B, C, E). Statistical analysis was performed by 2-way ANOVA (A-C, E) and TukeyHSD (A, B, E). Comparisons to wt are indicated in B, and comparisons to 30 min in E. ***, p < 0.001.

**Supplementary Figure S2.**
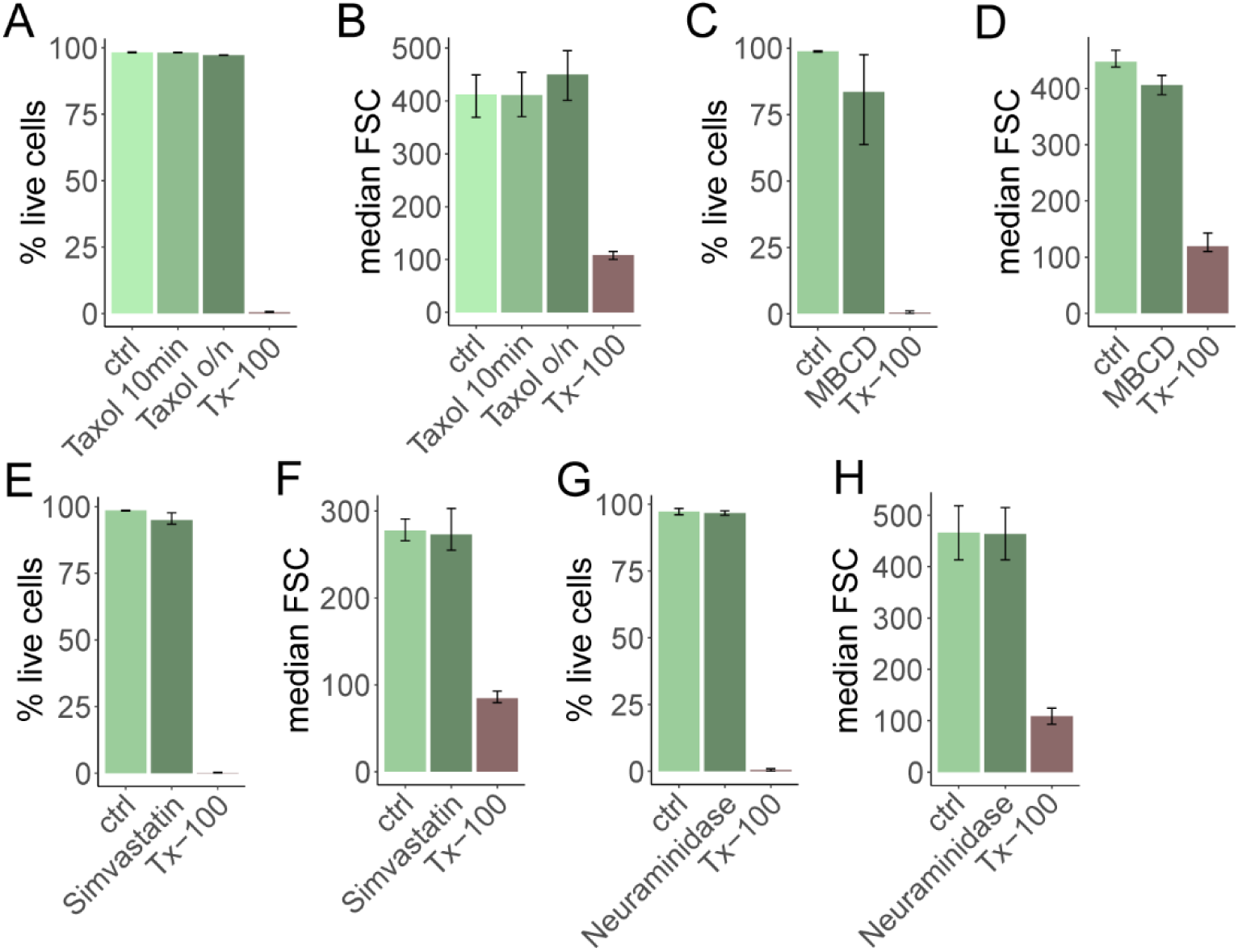
K562 viability following different cell surface manipulations. Viability as assessed by staining with propidium iodide (PI), and relative cell size measurements based on median forward scatter (FSC) following (A-B) treatment with Taxol for 10 min (as a control) or overnight (o/n) for mitotic arrest, (C-D) treatment with MBCD for 30 min or (E-F) with Simvastatin for 48 h for reduction of cholesterol levels, or (G-H) treatment with Neuraminidase for 1 h to trim the cell surface glycocalyx. For live/dead stainings, permeabilization with 0.1% Triton-X-100 (Tx-100) was included as a positive control. Data is plotted as mean + range of 3 replicates (A-F) or mean + sd of ≥ 4 replicates (G-H).

**Supplementary Figure S3.**
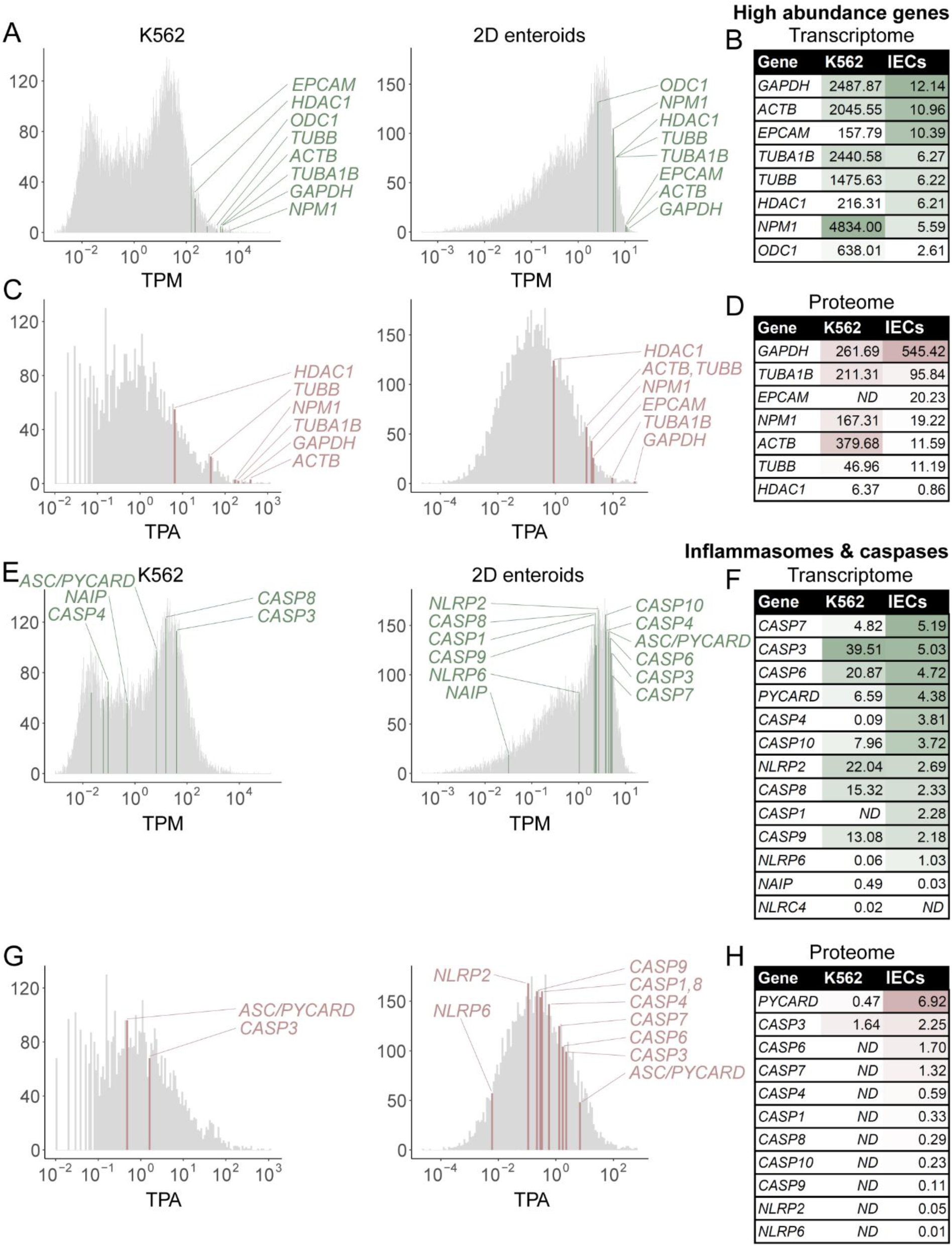
Transcriptome and proteome analysis of K562 cells at steady-state. Transcriptome and proteome analysis of K562 cells at steady-state and comparison to differentiated human 2D enteroid-derived IEC monolayers (data from (33)). (A, E) Transcripts with TPM (transcript per kilobase million) values >0 are plotted in a histogram, bins containing (A) well-known high abundance transcripts or (E) transcripts for inflammasome components and caspases are highlighted. (B, F) Lists of transcripts highlighted in A, E. For K562 cells, the mean of 4 replicates is depicted, while the mean of 3 replicates for independent enteroid lines derived from 2 patients is shown for IECs. (C, G) Identified proteins (≥1 unique + razor peptide) with TPA (total protein amount in fmol/µg total protein) values >0 are plotted in a histogram, bins containing (C) high abundance proteins or (G) inflammasome components or caspases are highlighted. (D, H) Lists proteins highlighted in C, G. The mean of 3 (IECs) or 4 (K562) replicates is depicted.

**Supplementary Figure S4.**
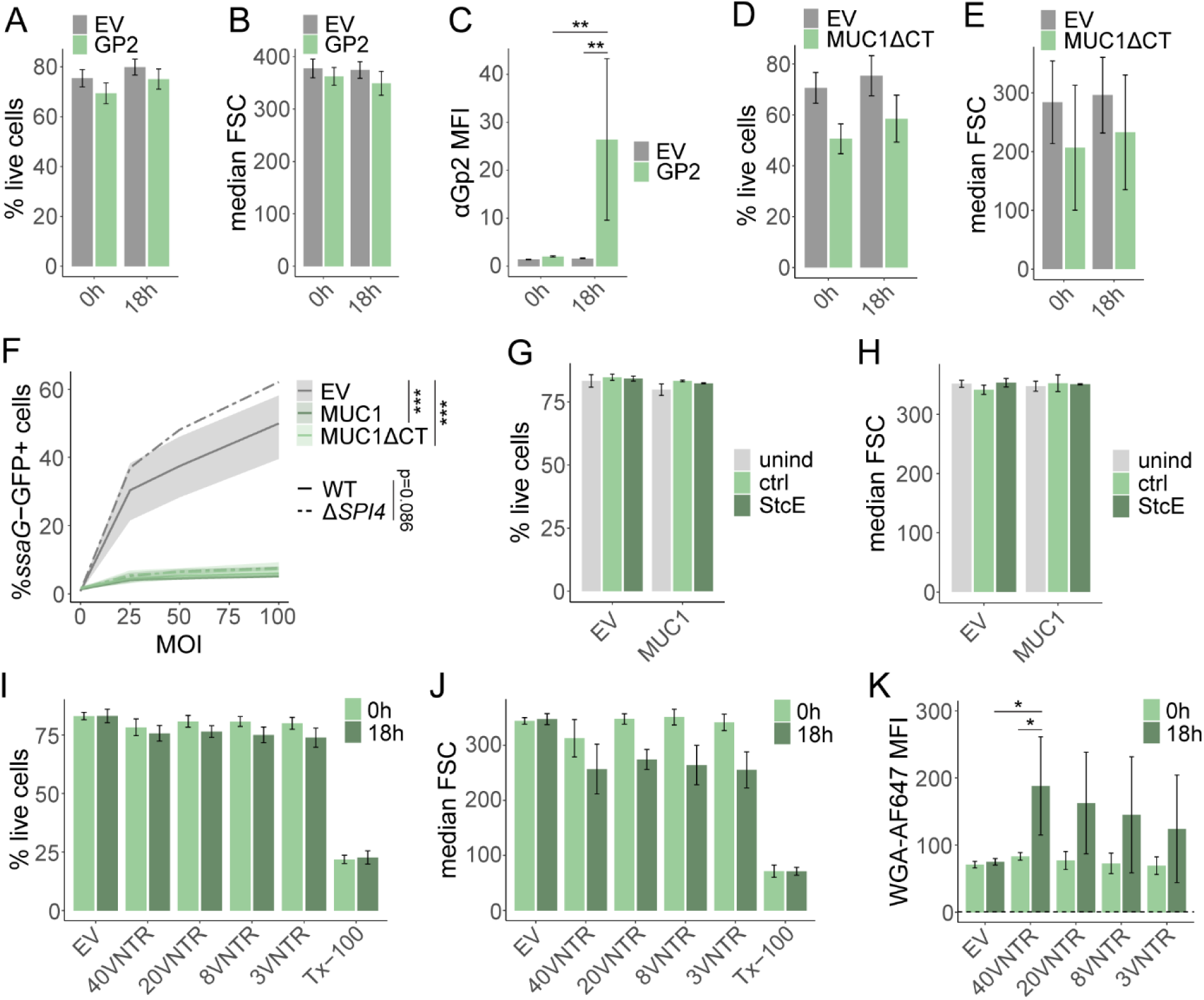
Additional data on cell surface manipulations in K562 cells. (A-B) Viability as assessed by PI staining (A) and relative cell size measurements based on median FSC (B) following transfection with pMEP4-GP2. Data is plotted as mean + sd of 8 replicates. (C) Staining with respective primary and secondary antibodies listed in Table S5, followed by flow cytometry confirms successful expression of GP2. Data is plotted as mean + sd of 4 replicates. (D-E) Viability as assessed by PI staining (D) and relative cell size measurements based on median FSC (E) following transfection with pMEP4-MUC1ΔCT. Data is plotted as mean + sd of 12 replicates. Time is indicated as hours post induction. (F) Both expression of full-length MUC1 and a truncated version lacking the cytoplasmic tail (MUC1ΔCT) in K562 cells results in massive reduction of *Salmonella* wt and Δ*SPI4* invasion. Data is plotted as mean + range of 3 replicates, except for the control (EV) infected with Δ*SPI4*, where only 1 replicate is plotted. A replicate corresponds to an individually transfected culture infected with 2 different inocula (mean for bacterial inocula). (G-H) Viability as assessed by PI staining (G) and relative cell size measurements based on median FSC (H) upon treatment of MUC1-transfected K562 cells with the mucinase StcE for 2 h. Data is plotted as mean + sd of 4 replicates. (I-J) K562 cells ectopically expressing HiBiT-tagged MUC1 versions with varying numbers of VNTRs, resulting in different lengths, display comparable cell viability as assessed by PI staining (I) and relative cell size measurements based on median FSC (J). (K) WGA staining of K562 cells expressing MUC1 versions of different lengths reveals that glycosylation levels are dependent on the number of VNTRs. Data is plotted as mean + sd of 6 replicates. Statistical analysis in C, F and K was performed by 2-way ANOVA and TukeyHSD. *, p < 0.05; **, p < 0.01; ***, p < 0.001; EV, empty vector; MFI, median fluorescence intensity.

**Supplementary Figure S5.**
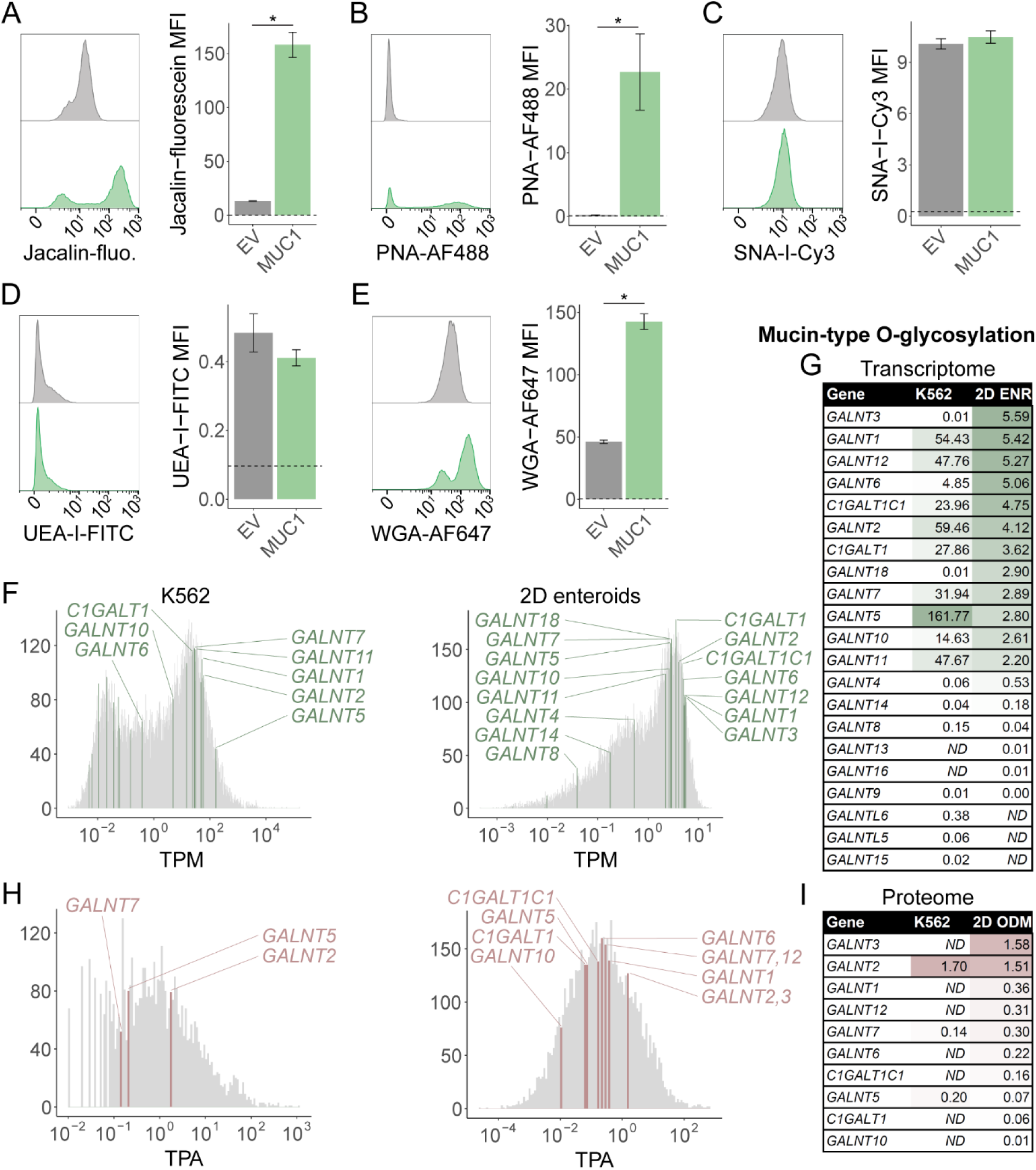
Analysis of mucin-type O-glycosylation in K562 cells. (A-E) MUC1-transfected K562 cells were stained with (A) Jacalin, (B) PNA, (C) SNA-I, (F) UEA-I, or (G) WGA to assess potential increases in cell surface glycosylation and gain insight into glycosylation patterns. Jacalin (A), PNA (B) and WGA stainings (E) show a robust increase upon ectopic expression of MUC1. Data is plotted as mean + sd of 4 replicates. Dashed lines indicate the mean of all unstained samples. Statistical analysis was performed by Mann-Whitney U test. *, p < 0.05. (F-I) Transcriptome (F-G) and proteome (H-I) analysis of glycosyltransferases involved in the first 2 steps of mucin-type O-glycosylation towards core 1 structures in differentiated human IECs (data from (33)) and K562 cells. (F) Transcripts with TPM > 0 are plotted in a histogram and bins containing transcripts for selected glycosyltransferases are highlighted. (G) List of TPM values for transcripts highlighted in F. The mean of 4 replicates is depicted for K562, while the mean of 3 replicates for independent human enteroid lines derived from 2 patients is shown for IECs. (H) Identified proteins (≥1 unique + razor peptide) with TPA > 0 are plotted in a histogram, bins containing selected glycosyltransferases are highlighted. (I) List of proteins highlighted in H. The mean of 3 (IECs) or 4 (K562) replicates is depicted. MFI, median fluorescence intensity; EV, empty vector.

**Supplementary Figure S6.**
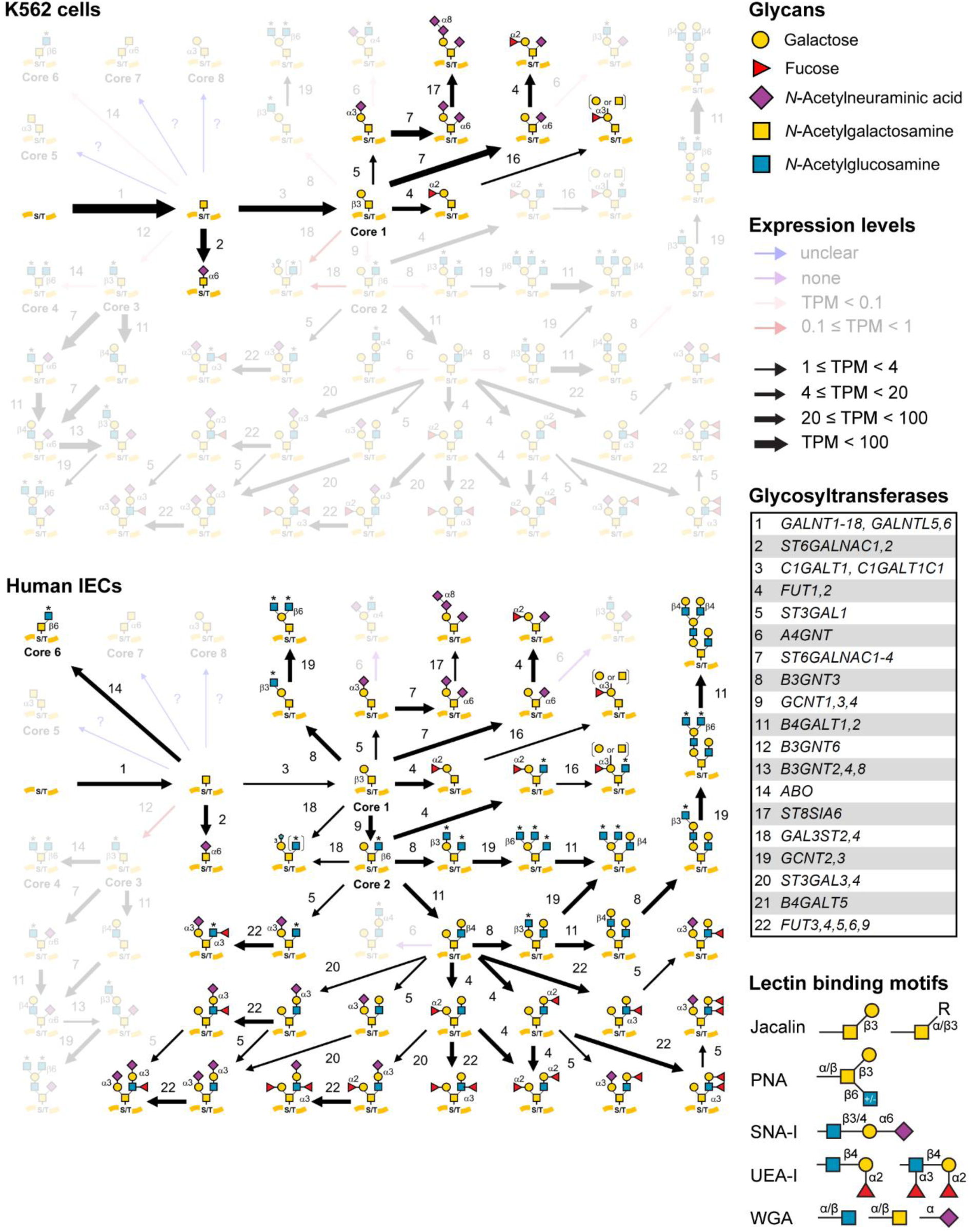
Predicted mucin-type O-glycan structures in K562 and human IECs. Transcriptome data for glycosyltransferases obtained from K562 cells or differentiated human IECs (data from (33)) were employed to predict mucin-type O-glycan structures using the Human GlycoMaple tool available at the GlyCosmos portal (https://glycosmos.org/glycomaple/Human) (69). The predictions reveal a higher diversity of O-glycan structures in human IECs, but indicate that glycan structures generated by K562 cells are present also in IECs. Pathways including at least one step that is catalyzed by a glycosyltransferase expressed at < 1 TPM are considered to have a minor contribution in mucin-type O-glycosylation and are depicted with decreased opacity. A list of all glycosyltransferases considered in the analysis (numbered 1-22) and relevant lectin binding motifs (according to (67)) are included.

**Table S1.** Strains and plasmids used in this study.

| Strain | Genotype | Reference |
| --- | --- | --- |
| *S. Tm* wt | SL1344, wt (SB300, Sm<sup>R</sup>) | (92) |
| *S. Tm* $\Delta invG$ | SL1344, $\Delta invG$ (SB161, Sm<sup>R</sup>) | (53) |
| *S. Tm* $\Delta sipA$ | SL1344, $\Delta sipA$ (M714, Sm<sup>R</sup>) | (102) |
| *S. Tm* $\Delta sopBEE2$ | SL1344, $\Delta sopB$, *sopE::aphT*, *sopE2::tetR* (M516, Sm<sup>R</sup>, Kan<sup>R</sup>, Tet<sup>R</sup>) | (103) |
| *S. Tm* $\Delta 4$ | SL1344, $\Delta sipA$, $\Delta sopB$, $\Delta sopE$, $\Delta sopE2$ (M566, Sm<sup>R</sup>) | (54) |
| *S. Tm* $\Delta SPI4$ | SL1344, $\Delta SPI4$ (Sm<sup>R</sup>, Kan<sup>R</sup>) | (28) |
| *S. Tm* $\Delta fimH$ | SL1344, $\Delta fimH$ (Sm<sup>R</sup>, Kan<sup>R</sup>) | This study |
| *S. Tm* $\Delta motA$ | SL1344, $\Delta motA$ (Sm<sup>R</sup>, Kan<sup>R</sup>) | (79) |
| *S. Tm* $\Delta fljBfliC$ | SL1344, $\Delta fljBfliC$ (Sm<sup>R</sup>, Kan<sup>R</sup>) | (104) |
| *S. Tm* $\Delta fljBfliC \Delta SPI4$ | SL1344, $\Delta fljBfliC \Delta SPI4$ (Sm<sup>R</sup>, Kan<sup>R</sup>) | (4) |
| *E. coli* Dh10b | Cloning strain used for amplification of pMEP4 vectors | - |
| Plasmid | Description | Reference |
| pM975 | *pssaG*-GFP (SPI2-dependent, vacuolar); Amp<sup>R</sup> | (47, 48) |
| *puhpT*-GFP | *puhpT*-GFP (cytosolic); Amp<sup>R</sup> | Related to *puhpT*-mCh in (49) |
| pFPV-mCherry | *rpsM*-mCherry (constitutive); Amp<sup>R</sup> | (52) |
| pMEP4-GP2 | Ectopic expression of GP2 in K562 cells; Amp<sup>R</sup>, Hyg<sup>R</sup> | This study |
| pMEP4-MUC1 | Ectopic expression of MUC1 in K562 cells; Amp<sup>R</sup>, Hyg<sup>R</sup> | This study |
| pMEP4-MUC1ΔCT | Ectopic expression of MUC1 without cytoplasmic tail in K562 cells; Amp<sup>R</sup>, Hyg<sup>R</sup> | This study |
| pMEP4-HiBiT-MUC1ΔCT (40 VNTR) | Ectopic expression of HiBiT-tagged MUC1ΔCT with full length extracellular domain (40 VNTR) in K562 cells; Amp<sup>R</sup>, Hyg<sup>R</sup> | This study |
| pMEP4-HiBiT-MUC1ΔCT-20VNTR | Ectopic expression of HiBiT-tagged MUC1ΔCT with extracellular domain comprising 20 VNTR in K562 cells; Amp<sup>R</sup>, Hyg<sup>R</sup> | This study |
| pMEP4-HiBiT-MUC1ΔCT-8VNTR | Ectopic expression of HiBiT-tagged MUC1ΔCT with extracellular domain comprising 8 VNTR in K562 cells; Amp<sup>R</sup>, Hyg<sup>R</sup> | This study |
| pMEP4-HiBiT-MUC1ΔCT-3VNTR | Ectopic expression of HiBiT-tagged MUC1ΔCT with extracellular domain comprising 3 VNTR in K562 cells; Amp<sup>R</sup>, Hyg<sup>R</sup> | This study |
| CMV-EBNA1 | Improved expression of pMEP4-encoded sequences in K562 cells; Amp<sup>R</sup>, Hyg<sup>R</sup> | (105) |

Sm, Streptomycin; Kan, Kanamycin; Tet, Tetracyclin; Amp, Ampicilin; Hyg, Hygromycin; mCh, mCherry.

**Table S2.**
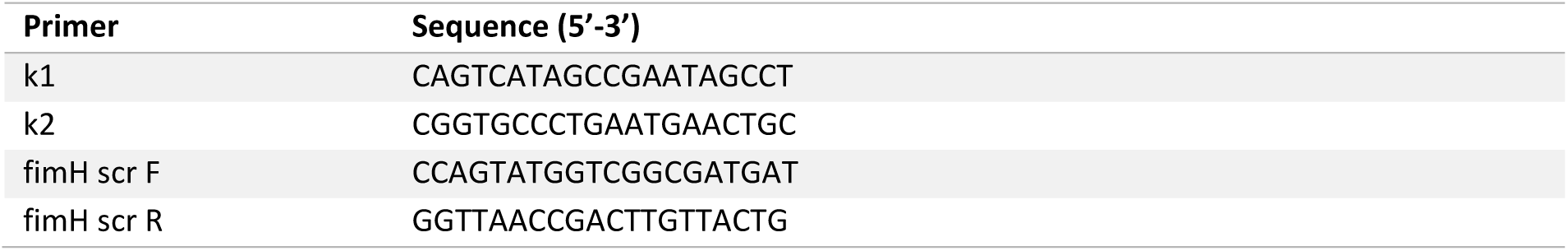
Primers used in this study.

**Table S3.**
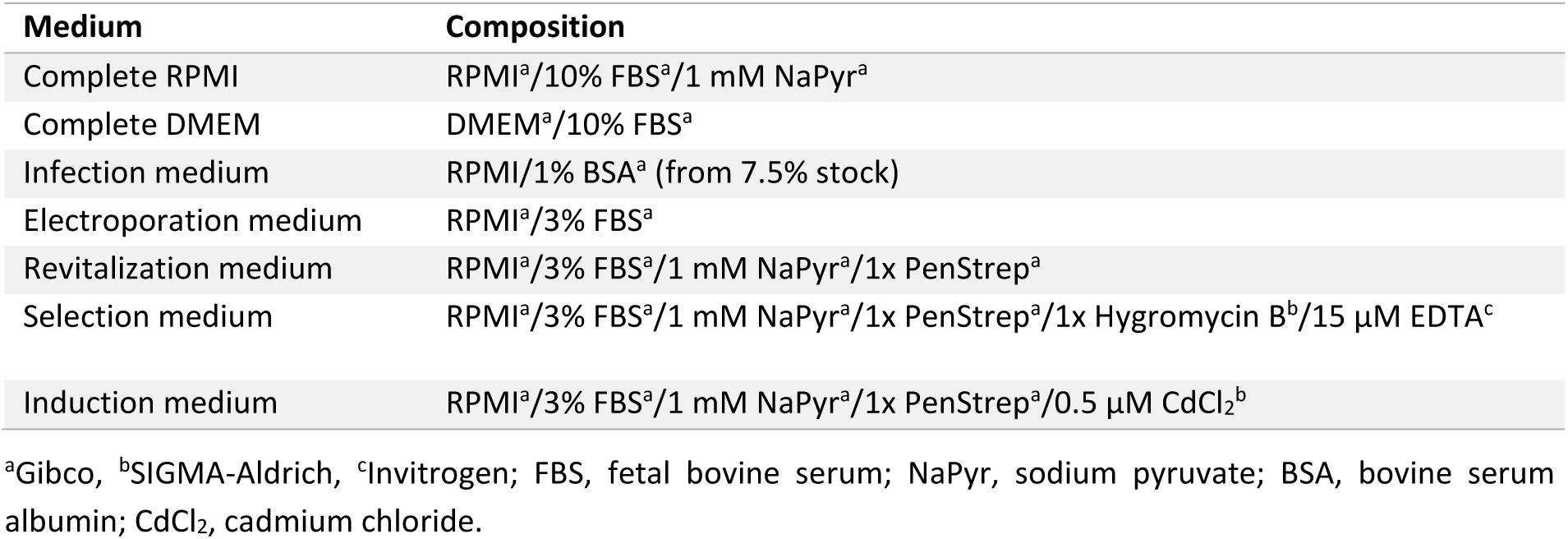
Overview of cell culture media.

**Table S4.**
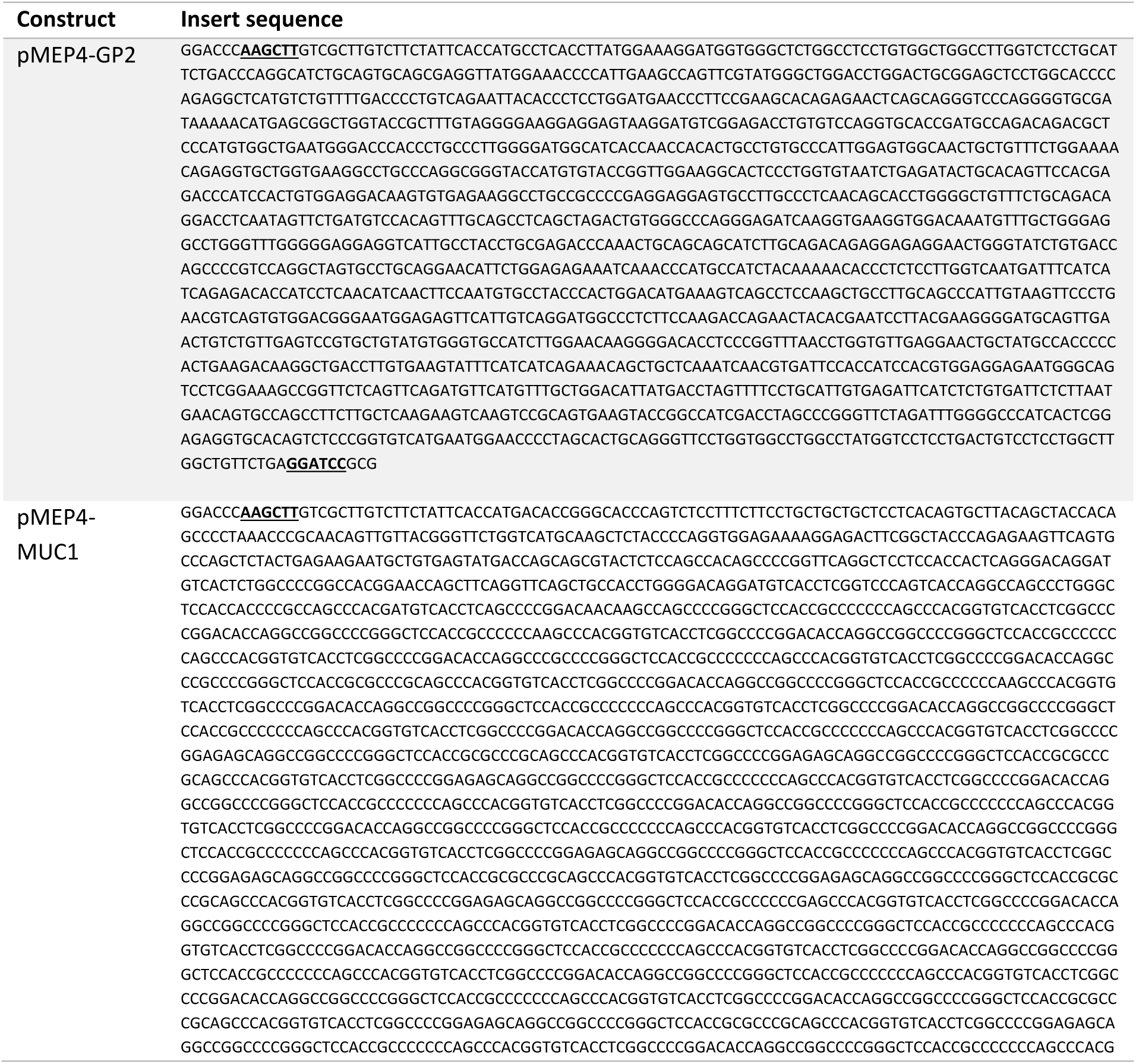
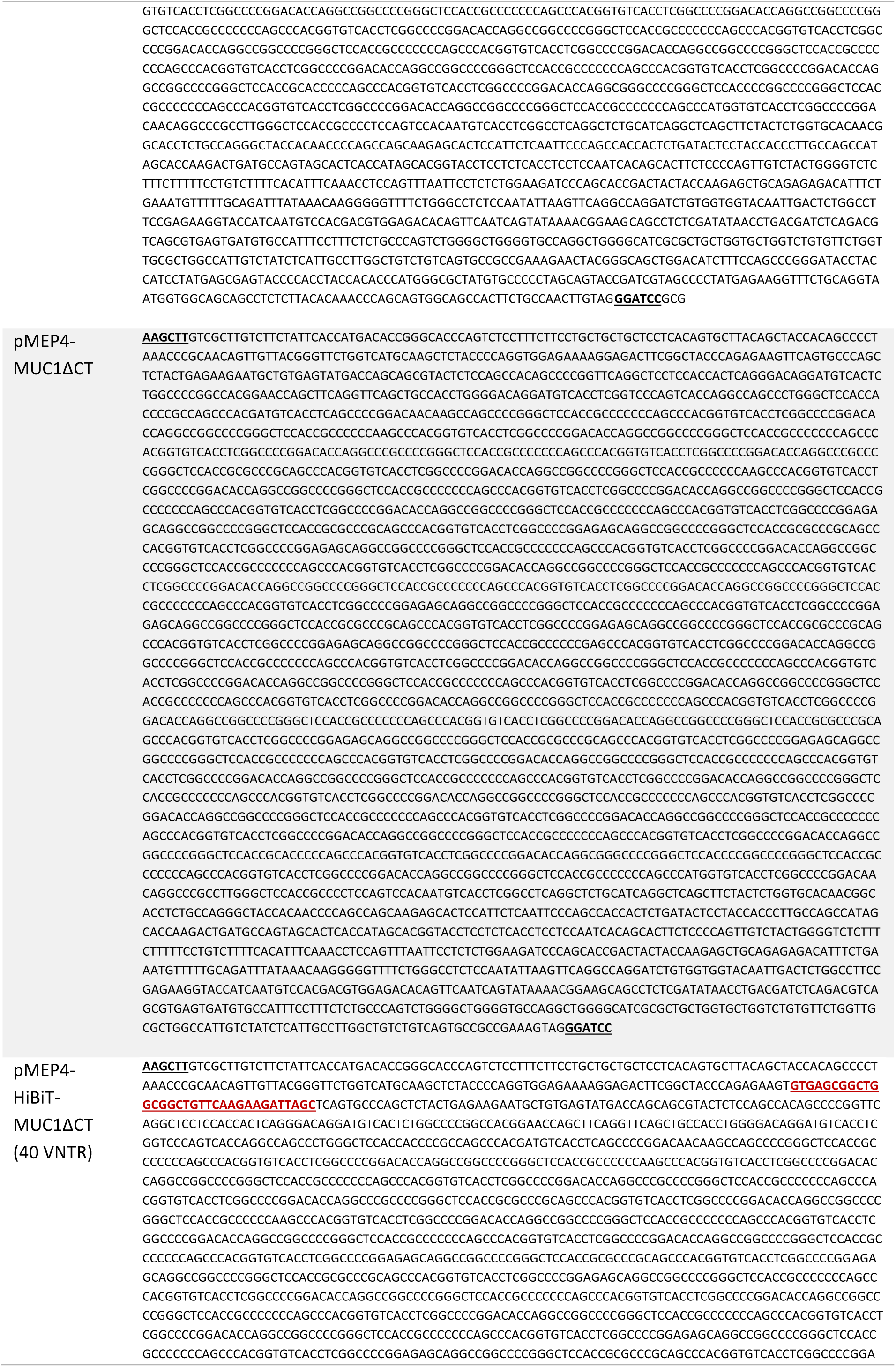
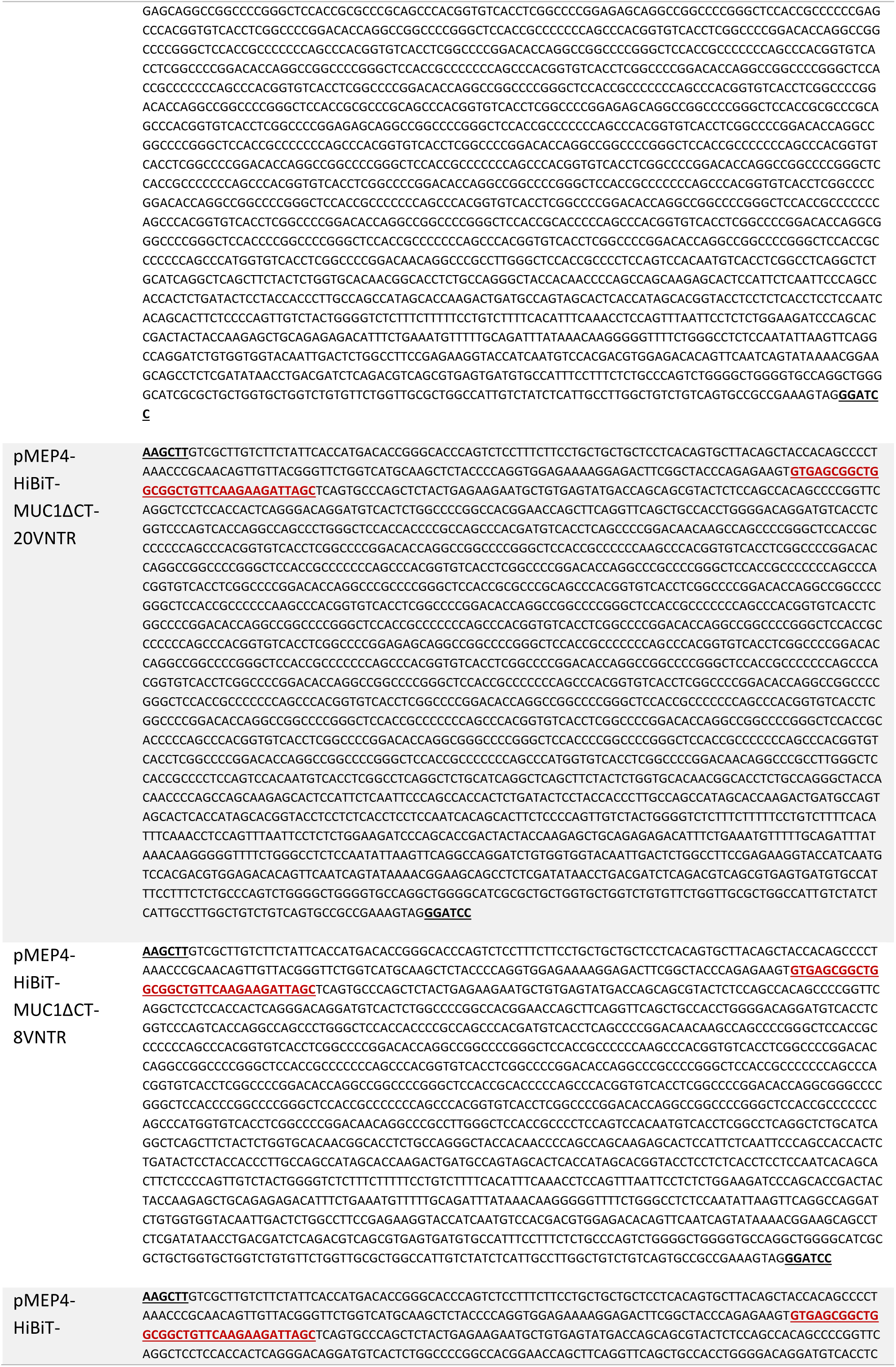
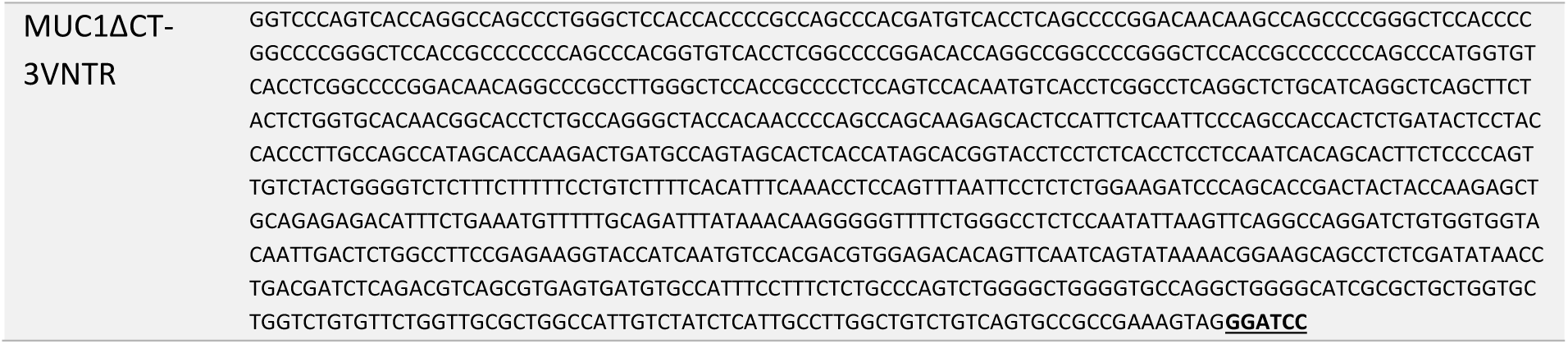
Insert sequences for pMEP4 vectors used in this study. HindIII and BamHI restriction sites as well as HiBiT-tag sequences are indicated.

**Table S5.**
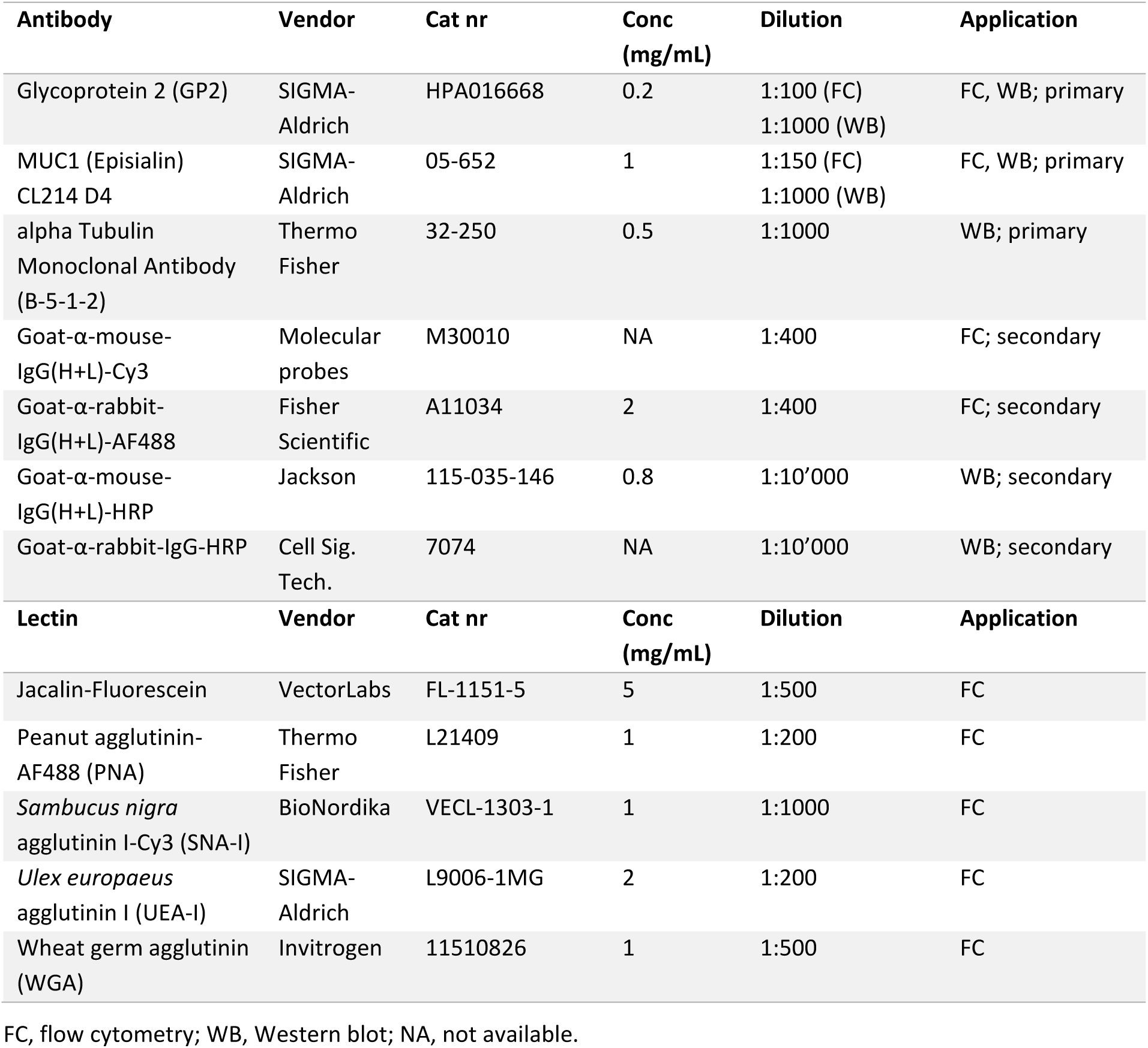
Antibodies and lectins used in this study.

## Notes

### Competing Interest Statement

The authors have declared no competing interest.

